# HIF-1α integrates metabolic and immunoregulatory programs in RORγt⁺ regulatory T cells during intestinal inflammation

**DOI:** 10.64898/2026.08.11.744213

**Authors:** Marcella Cipelli, Eloísa M. da Silva, Luísa Menezes-Silva, Barbara N. Padovani, Mariana A. Amaral, Laís C. Paredes, Bruno G. Nunes, Victor Y. Yariwake, José Arimatéia O. N. Neto, Natalia N. Bos, Anthony G. da Silveira, João Vinicius H. da Silva, Raquel S. Vieira, Suemy M. Yamada, Luis Felipe S. Moreira, Benedito Matheus dos Santos, Aline Ignacio, Maria Fernanda Forni, Orestes Foresto-Neto, Jefferson Antonio Leite, Marco Aurélio R. Vinolo, Dennyson Leandro M. da Fonseca, Sandra Marcia Muxel, Matthias Lochner, Vinicius Andrade-Oliveira, Niels O. S. Camara

## Abstract

Regulatory T (Treg) cells expressing RORγt accumulate in the intestinal mucosa, yet the signals that determine whether they remain suppressive or acquire inflammatory features are incompletely defined. We first reanalyzed human ileal single-cell data and identified Crohn’s disease–enriched FOXP3⁺ states in which RORC, HIF1A, hypoxia-responsive, inflammatory, and metabolic programs converged. We then deleted Hif1a in RORγt-expressing cells and tested acute DSS colitis, T cell transfer colitis, and azoxymethane/DSS-induced colitis-associated colorectal cancer (CAC). ΔHif1a mice were protected in all three settings. In lymphopenic recipients given the same pathogenic naïve T cells, changing only the genotype of the cotransferred Treg population enhanced protection, linking the phenotype to regulatory-cell function *in vivo*. Reanalysis of mouse colonic Treg single-cell ATAC-seq nominated suppressive and mitochondrial programs for cell-intrinsic testing during low HIF1-α expression. ΔHif1a RORγt⁺ Treg produced more IL-10 and less IL-17A and IFN-γ, limited responder-cell proliferation, contained fewer dysfunctional and mitochondrial-reactive-oxygen-species-high mitochondria, favored fusion-associated transcription, and displayed greater basal and maximal oxygen consumption and reserve capacity. During CAC, HIF-1α loss blunted inflammatory RORγt⁺ Treg accumulation and reduced tumor burden. Human trajectory and gene-regulatory-network analyses further predicted that HIF1A perturbation would oppose selected disease-associated branches. Together, these findings identify HIF-1α as a context-dependent checkpoint that connects hypoxia-responsive transcription to mitochondrial fitness and inflammatory plasticity in intestinal RORγt⁺ Treg.

## INTRODUCTION

At mucosal surfaces, CD4⁺ T cells must defend the barrier without converting a sustained response into tissue injury. RORγt-expressing T helper 17 (Th17) cells contribute to barrier defense but can drive inflammatory bowel disease (IBD) when dysregulated. Foxp3⁺ Treg restrain this response, and a microbiota-responsive subset that coexpresses RORγt is particularly abundant in the intestine^1,4–8^. These RORγt⁺ Treg are not functionally uniform: depending on tissue and inflammatory context, they can sustain tolerance and IL-10 production or acquire inflammatory cytokines while retaining regulatory markers^9–25^.

This plasticity is clinically relevant since RORγt⁺ or IL-17-producing FOXP3⁺ cells accumulate in human IBD and colorectal cancer, where local signals can alter both their suppressive and inflammatory functions^10,11,13,16,21,26,27,44^. Single-cell analysis of ileal CD4⁺ T cells recently identified Crohn’s disease–enriched FOXP3⁺ populations that respond to TNF and exhibit impaired suppression^62^. These observations define a disease-associated state, but they do not explain how the hypoxic and metabolically constrained intestinal environment helps establish it.

Cellular metabolism is an active determinant of T cell fate. mTOR integrates nutrient, cytokine, and antigen-receptor signals, whereas HIF-1α coordinates adaptation to low oxygen and inflammatory stress^28–42^. In conventional CD4⁺ T cells, HIF-1α promotes glycolysis and Th17 differentiation and can antagonize Foxp3 stability^40,41^. Intestinal RORγt⁺ Treg reside where hypoxia, microbiota-derived metabolites, and inflammatory mediators converge. We therefore reasoned that HIF-1α might control not simply the abundance of this population, but the functional and mitochondrial state it adopts during inflammation.

Here, we used the Crohn’s disease dataset GSE209832 to position HIF1A within human RORγt-associated FOXP3⁺ states, then tested its function genetically in acute DSS colitis, T cell transfer colitis, and AOM/DSS-induced CAC. The transfer model was used to standardize the lymphopenic recipient, pathogenic T cell inoculum, and myeloid environment while varying the cotransferred Treg. Mouse colonic Treg chromatin accessibility was analyzed before cell-intrinsic experiments to nominate regulatory and mitochondrial outputs, which were then evaluated by differentiation, suppression, mitochondrial-dye, RT–qPCR, and extracellular-flux assays. Finally, human trajectory inference and *in silico* HIF1A perturbation were used to ask whether the mouse mechanism mapped onto specific disease-associated regulatory branches in humans.

## RESULTS

### Crohn’s disease–associated Treg states are enriched for RORγt, HIF1A, and hypoxia-responsive programs

We began by asking whether HIF1A was associated with the pro-inflammatory regulatory states identified in Crohn’s disease. We reanalyzed scRNA-seq data from ileal CD4⁺ T cells from healthy donors and patients with Crohn’s disease (GSE209832)^62^. Reclustering the FOXP3⁺ compartment resolved seven transcriptional states (clusters 0–6; Fig. 1A). RORγt-high FOXP3⁺ cells occupied partially distinct regions of the embedding and were more prominent in Crohn’s disease samples (Fig. 1B), focusing the subsequent analysis on disease-associated RORγt⁺ Treg-like states.

**Figure 1.**
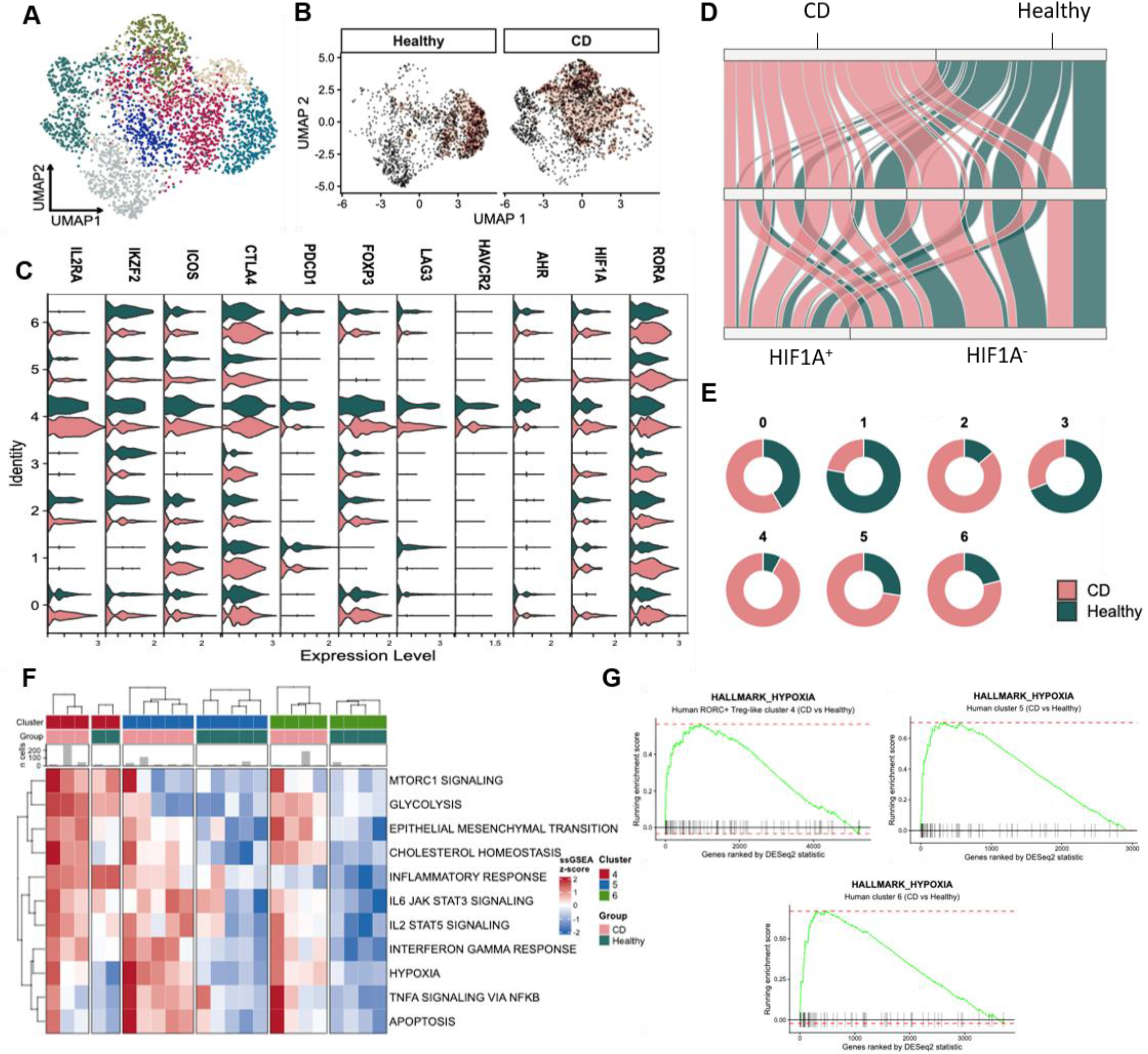
Crohn’s disease–associated Treg cell states are enriched for RORγt, HIF1A, and hypoxia-responsive programs. Reanalysis of ileal CD4⁺ T cell scRNA-seq data from healthy donors and patients with Crohn’s disease (CD; GSE209832). (A) UMAP of FOXP3⁺ cells colored by clusters 0–6. (B) Density of RORγt-high FOXP3⁺ cells in healthy and CD samples. (C) Split violin plots showing normalized expression of regulatory, inhibitory, and environment-responsive genes across clusters and conditions. (D) Alluvial diagram linking condition, cluster identity, and detectable HIF1A transcript (HIF1A⁺ versus HIF1A⁻). (E) Proportion of healthy and CD cells in each cluster. (F) Heatmap of standardized ssGSEA scores for Hallmark pathways across patient– cluster combinations; annotation bars indicate cluster and disease group. (G) Preranked GSEA plots for HALLMARK_HYPOXIA in cluster 4 (RORC⁺ Treg-like), cluster 5, and cluster 6, comparing CD with healthy samples.

Regulatory and inhibitory genes (IL2RA, IKZF2, ICOS, CTLA4, PDCD1, FOXP3, LAG3, and HAVCR2) and environment-responsive factors (AHR, HIF1A, RORA, and RORC) varied across clusters and conditions (Fig. 1C). Linking disease status, cluster identity, and detectable HIF1A transcripts showed that HIF1A-positive cells were distributed nonuniformly across healthy- and Crohn’s disease–associated states (Fig. 1D). Clusters 2, 4, 5, and 6 were enriched for Crohn’s disease cells, whereas clusters 1 and 3 were predominantly healthy (Fig. 1E). Thus, HIF1A expression did not mark every FOXP3⁺ cell; it was preferentially associated with selected disease-enriched states.

We next asked which programs accompanied this distribution. Sample-aware ssGSEA separated patient–cluster combinations by mTORC1 signaling, glycolysis, inflammatory response, IL-6–JAK–STAT3, IL-2–STAT5, interferon-γ, hypoxia, TNF–NF-κB, and apoptosis (Fig. 1F). Independently, preranked GSEA showed positive enrichment of the hypoxia program in Crohn’s disease relative to healthy cells within the RORC⁺ Treg-like cluster 4 and clusters 5 and 6 (Fig. 1G). The convergence of RORC, HIF1A detection, and hypoxia/inflammatory activity in these human states provided the rationale to test whether HIF-1α actively controls intestinal RORγt⁺ Treg function.

### HIF-1α deletion in RORγt-expressing cells attenuates acute DSS colitis and preserves a suppressive RORγt⁺ Treg phenotype

We first established where HIF-1α was most evident among murine CD4⁺ T cell subsets. RORγt⁺Foxp3⁺ Treg showed greater HIF-1α staining than Foxp3⁺RORγt⁻ conventional Treg or Foxp3⁻RORγt⁺ Th17 cells (Fig. 2A), and the difference was greatest in the colonic lamina propria (CLP) compared with spleen (SPL), mesenteric lymph nodes (MLN), and peripheral lymph nodes (PLN) (Fig. 2B). We therefore crossed Rorc^Cre^ mice with Hif1a^flox/flox^ mice to generate ΔHif1a animals, in which HIF-1α is deleted in cells with a RORγt expression. Thymic CD4/CD8 composition and CD5 and CD69 profiles were broadly preserved (Fig. S1), showing that Hif1a deletion did not compromise normal lymphocyte development.

**Figure 2.**
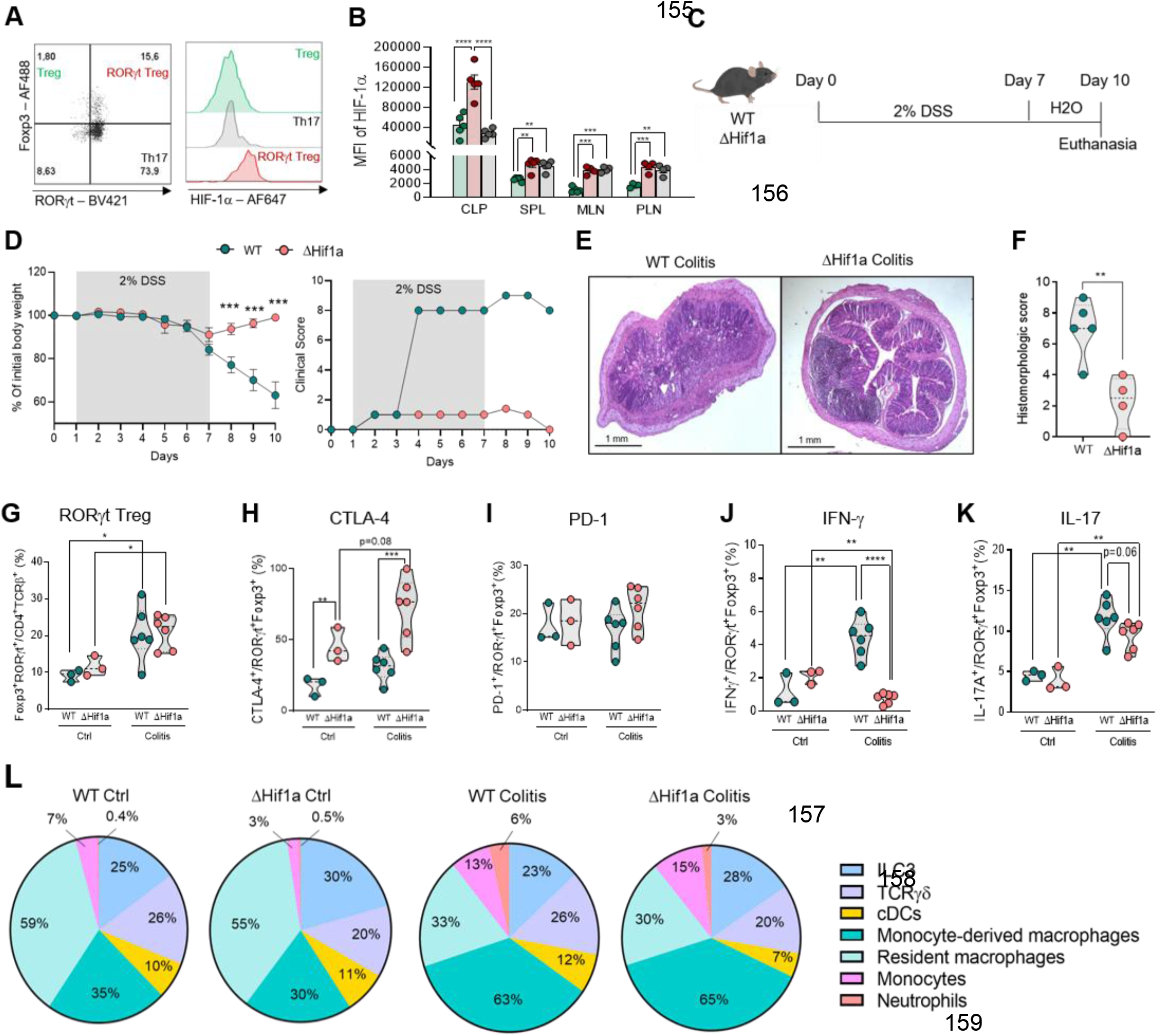
HIF-1α deletion in RORγt-expressing cells attenuates acute DSS colitis and preserves a suppressive RORγt⁺ Treg phenotype. (A) Representative Foxp3/RORγt gating of conventional Treg, RORγt⁺ Treg, and Th17 cells with overlaid HIF-1α histograms. (B) HIF-1α mean fluorescence intensity across the indicated CD4⁺ subsets from CLP, SPL, MLN, and PLN. (C) Acute-colitis design: 2% DSS on days 0–7, water on days 7–10, and euthanasia on day 10. (D) Body-weight change and clinical score. (E) Representative H&E-stained colon sections. (F) Histomorphologic score. (G) Frequency of CLP Foxp3⁺RORγt⁺ Treg. (H–K) CTLA-4 (H), PD-1 (I), IFN-γ (J), and IL-17A (K) in CLP RORγt⁺ Treg. (L) Relative CLP composition of ILC3s, TCRγδ cells, conventional dendritic cells, monocyte-derived macrophages, resident macrophages, monocytes, and neutrophils. Points represent mice; bars or violins show mean ± SEM.

WT and ΔHif1a mice next received 2% DSS for seven days followed by water through day 10 (Fig. 2C). Compared with WT mice, ΔHif1a mice lost less weight, developed lower clinical scores, retained epithelial architecture, and had lower histomorphologic scores (Fig. 2D–F). RORγt⁺ Treg frequencies increased during colitis in both genotypes (Fig. 2G), showing that protection was not caused by elimination of this population. Instead, ΔHif1a RORγt⁺ Treg expressed more CTLA-4 at baseline and during colitis, while PD-1 frequencies were not significantly changed (Fig. 2H,I). IFN-γ production was markedly reduced during colitis, and IL-17A was numerically lower (Fig. 2J,K), consistent with a shift away from an inflammatory regulatory phenotype.

Complementary profiling revealed a tissue-specific redistribution rather than generalized loss of the regulatory compartment. In CLP, Treg frequencies declined numerically during DSS colitis, whereas Th17 frequencies increased in both genotypes, without a genotype-specific Th17 increase that could account for the ΔHif1a phenotype (Fig. S2B). In spleen, total Foxp3⁺ Treg rose numerically during colitis in both genotypes, whereas ΔHif1a mice showed a pronounced expansion of Foxp3⁺RORγt⁺ Treg relative to WT mice with colitis; splenic Th17 frequencies did not show a corresponding genotype-dependent expansion (Fig. S2C). Despite this accumulation, ΔHif1a splenic RORγt⁺ Treg expressed more CTLA-4 and CD44 at baseline and during colitis, less PD-1 during colitis, and markedly less IFN-γ and IL-17A (Fig. S2D,E). Colon tissue IL-10 was significantly increased in ΔHif1a mice with colitis (Fig. S2F). Thus, HIF-1α deletion did not alter the frequency of extraintestinal RORγt⁺ Treg but reduced their inflammatory cytokine production and increased expression of a regulatory mediator in the affected tissue. Broad profiling of ILC3s, TCRγδ cells, conventional dendritic cells, monocytes, neutrophils, and macrophage subsets did not reveal a compositional change proportional to the degree of protection (Fig. 2L; gating in Fig. S3), although Rorc^Cre^ also targets other RORγt-lineage cells.

### ΔHif1a Treg provide enhanced protection in T cell transfer colitis

To isolate the *in vivo* contribution of the regulatory population from broader effects of RorcCre-driven deletion, we used the T cell transfer colitis model. Rag1⁻/⁻ recipients lack endogenous T and B lymphocytes but retain an otherwise comparable recipient myeloid compartment. Every cotransfer group received the same WT naïve CD4⁺ pathogenic inoculum; the experimental variable was whether the accompanying CD4⁺CD25^high^ Treg came from WT or ΔHif1a donors. Recipients received naïve cells alone or with Treg at a 4:1 naïve:Treg ratio and were followed for five weeks (Fig. 3A)^45^. Naïve cells alone caused progressive weight loss, colon shortening, splenomegaly, and severe colonic inflammation. WT Treg provided partial protection, whereas ΔHif1a Treg sustained body weight more effectively, preserved colon length, limited splenomegaly, and reduced mucosal injury (Fig. 3B–F).

**Figure 3.**
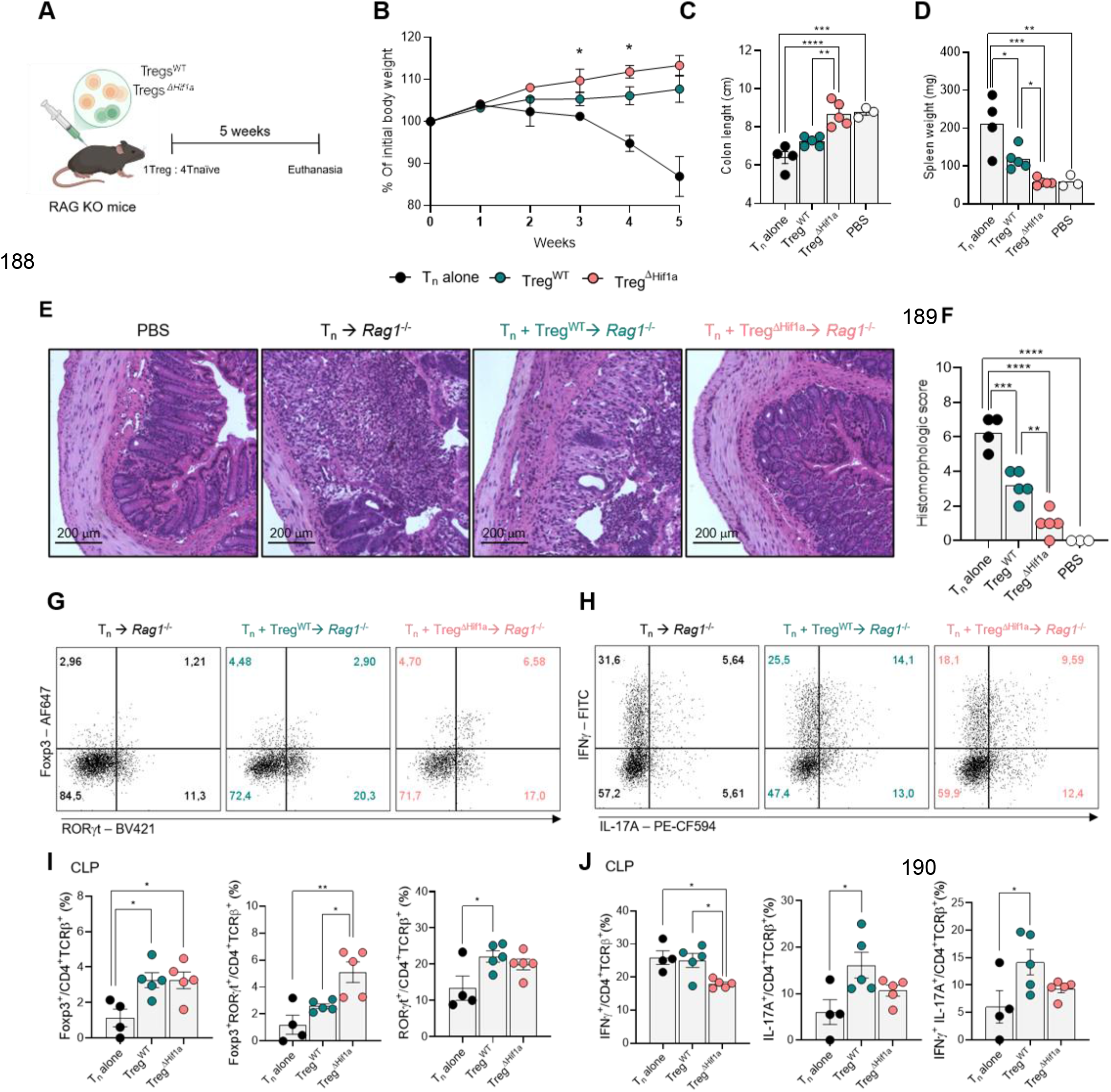
ΔHif1a Treg provide enhanced protection in T cell transfer colitis. (A) Experimental design. Rag1⁻/⁻ recipients received the same WT naïve CD4⁺ T cell inoculum alone or together with CD4⁺CD25high WT or ΔHif1a Treg at a 4:1 naïve:Treg ratio and were evaluated after five weeks. (B) Body-weight change. (C) Colon length. (D) Spleen weight. (E) Representative H&E-stained colon sections from PBS, naïve T cell alone, naïve plus WT Treg, and naïve plus ΔHif1a Treg groups. Scale bars, 200 µm. (F) Histomorphologic score. (G) Representative Foxp3/RORγt profiles in CLP CD4⁺ T cells. (H) Representative IFN-γ/IL-17A profiles. (I) Frequencies of Foxp3⁺ Treg, Foxp3⁺RORγt⁺ Treg, and RORγt⁺ CD4⁺ T cells in CLP. (J) Frequencies of IFN-γ⁺, IL-17A⁺, and IFN-γ⁺IL-17A⁺ CD4⁺ T cells in CLP. Points represent recipient mice; bars show mean ± SEM.

Flow cytometry of the CLP showed recovery of Foxp3⁺ Treg in both cotransfer groups and a larger Foxp3⁺RORγt⁺ fraction in recipients of ΔHif1a Treg (Fig. 3G,I). In the same mice, effector CD4⁺ T cells contained fewer IFN-γ⁺, IL-17A⁺, and IFN-γ⁺IL-17A⁺ cells than in recipients of WT Treg (Fig. 3H,J). Because the recipient genotype, lymphopenic setting, pathogenic T cell inoculum, and type of myeloid environment were standardized, the stronger protection can be attributed to the genotype of the transferred regulatory population within this model.

### HIF-1α loss enhances suppressive function and mitochondrial fitness *in vitro*

To prioritize cell-intrinsic pathways for functional testing, we first reanalyzed publicly available scATAC-seq data from mouse colonic Treg (GSM6697676)46. RORγt⁺ and RORγt⁻ Treg occupied related but distinguishable chromatin states while retaining gene activity at canonical regulatory loci (Fig. S4A–C). Relative to RORγt⁻ Treg, RORγt⁺ Treg showed a distinct accessibility profile at the Hif1a locus and higher gene activity scores for Ctla4, Icos, Lag3, Havcr2, and Il10, whereas Foxp3, Ikzf2, Il2ra, and Tigit were more prominent in the RORγt⁻ state (Fig. S4D, E). Metabolic gene activity was also redistributed across hypoxia-responsive, pyruvate-handling, mitochondrial-dynamics, and AHR-associated pathways (Fig. S4F). Consistent with these differences, chromVAR analysis showed greater HIF1A, AHR, NFATC2, STAT5/STAT5A, and MLXIP motif activity in RORγt⁺ Treg, whereas NR4A2, LXR, ESRRG, and PPAR motifs predominated in RORγt⁻ Treg (Fig. S4G, H).

We next stratified the RORγt⁺ compartment according to HIF1A motif deviation scores. HIF1A motif-high and motif-low states differed in accessible peaks and segregated regulatory genes into distinct modules. HIF1A motif-high cells showed greater Foxp3, Ikzf2, Lag3, and Il10 activity, whereas motif-low cells showed greater Il2ra, Ctla4, Tigit, Icos, and Havcr2 activity (Fig. S4I, J). Metabolic programs were similarly redistributed: the motif-low state included higher Cpt1a, Opa1, and Mfn2 activity, whereas the motif-high state favored a stress- and glycolysis-associated module containing Pdk3 (Fig. S4K). Because motif deviation reflects chromatin accessibility surrounding transcription factor motifs rather than HIF-1α protein abundance or activity, these observations were used to nominate regulatory function, cytokine production, mitochondrial dynamics, and oxidative metabolism for direct experimental evaluation following genetic HIF-1α deletion.

We generated RORγt⁺ Treg from naïve CD4⁺ T cells through four days of Treg polarization followed by IL-6-containing restimulation until day 8 (Fig. 4A)74. Compared with WT cultures, ΔHif1a Foxp3⁺RORγt⁺ cells expressed more PD-1 and showed a modest increase in CTLA-4, accompanied by increased IL-10 secretion and markedly reduced IL-17A production (Fig. 4B–E). We then assessed their ability to suppress conventional T cell proliferation. WT or ΔHif1a RORγt⁺ Treg were cultured with CellTrace-labeled CD4⁺ responder T cells activated with soluble anti-CD3 in the presence of Rag-deficient splenocytes. ΔHif1a RORγt⁺ Treg reduced responder-cell proliferation at Treg ratios ranging from 1:8 to 1:1, whereas suppression was comparable between genotypes at the lowest ratio tested, 1:16 (Fig. 4F). Thus, the less inflammatory cytokine profile of ΔHif1a RORγt⁺ Treg was accompanied by enhanced suppressive activity.

**Figure 4.**
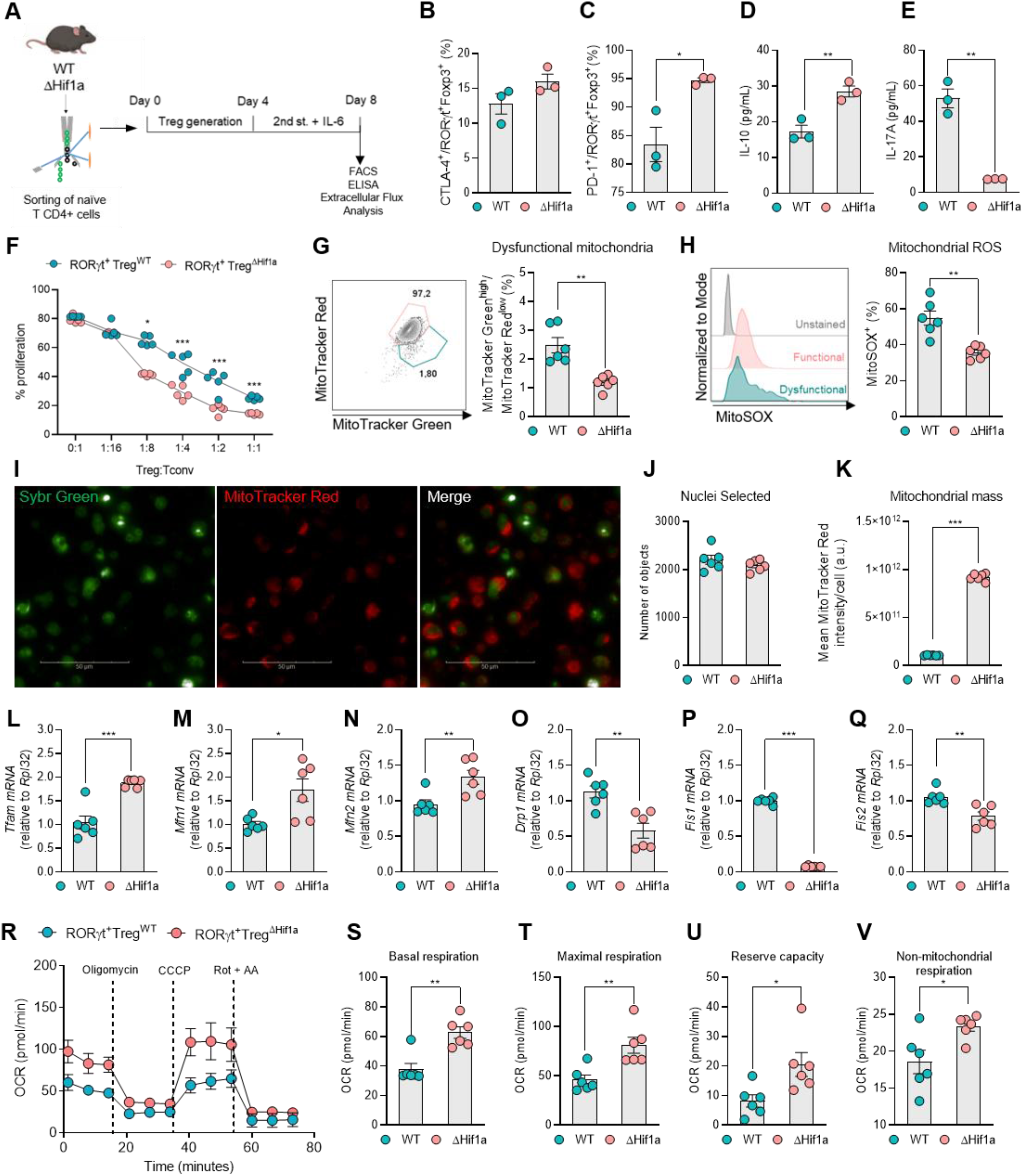
HIF-1α loss enhances suppressive function and mitochondrial fitness *in vitro*. (A) Experimental workflow showing naïve CD4⁺ T cell isolation, Treg polarization from days 0 to 4, IL-6-containing restimulation from days 4 to 8, and endpoint analyses. (B, C) Frequencies of CTLA-4⁺ (B) and PD-1⁺ (C) cells among Foxp3⁺RORγt⁺ Treg. (D, E) Concentrations of IL-10 (D) and IL-17A (E) in culture supernatants. (F) Proliferation of CellTrace Violet–labeled CD4⁺ responder T cells stimulated with soluble anti-CD3 in the presence of Rag-deficient splenocytes and cocultured with WT or ΔHif1a RORγt⁺ Treg at the indicated Treg ratios. (G) Representative MitoTracker Green/Red plot and frequency of MitoTracker Green^high^/MitoTracker Red^low^ cells within the dysfunctional-mitochondria gate. (H) Representative MitoSOX fluorescence histograms and frequency of MitoSOX⁺ cells. (I) Representative confocal images of SYBR Green– stained nuclei, MitoTracker Red–labeled mitochondria, and merged channels. Scale bar, 50 μm. (J) Number of nuclei selected for image analysis. (K) Mean MitoTracker Red fluorescence intensity per cell. (L–Q) Relative mRNA expression of Tfam (L), Mfn1 (M), Mfn2 (N), Drp1 (O), Fis1 (P), and Fis2 (Q). (R) Oxygen consumption rate (OCR) during the mitochondrial stress test, with the sequential addition of oligomycin, CCCP, and rotenone plus antimycin A (Rot + AA). (S– V) Basal respiration (S), maximal respiration (T), reserve capacity (U), and non-mitochondrial respiration (V). Points represent independent cultures; data are presented as mean ± SEM. *P < 0.05, **P < 0.01, and ***P < 0.001.

We next examined whether this functional phenotype was associated with changes in mitochondrial integrity. Flow cytometry showed that ΔHif1a cultures contained fewer MitoTracker Green^high/MitoTracker Red^low cells bearing dysfunctional mitochondria and exhibited lower MitoSOX-detectable mitochondrial reactive oxygen species (Fig. 4G, H). Confocal microscopy using SYBR Green to identify nuclei and MitoTracker Red to visualize mitochondria provided an independent assessment of the mitochondrial phenotype (Fig. 4I). Comparable numbers of nuclei were analyzed in WT and ΔHif1a cultures, excluding differences in cell representation during image quantification (Fig. 4J). In contrast, the mean MitoTracker Red fluorescence per cell was markedly increased in ΔHif1a cultures (Fig. 4K). Together with the flow-cytometric measurements, these imaging data indicate that Hif1a deletion is associated with a greater mitochondrial signal and improved mitochondrial integrity. RT–qPCR further revealed increased Tfam, Mfn1, and Mfn2 expression and reduced Drp1, Fis1, and Fis2 expression in ΔHif1a RORγt⁺ Treg, consistent with improved mitochondrial maintenance and a transcriptional shift toward fusion-associated rather than fission-associated dynamics (Fig. 4L–Q). During a mitochondrial stress test, ΔHif1a RORγt⁺ Treg maintained higher oxygen consumption and displayed increased basal respiration, maximal respiration, and reserve capacity, together with a small increase in non-mitochondrial oxygen consumption (Fig. 4R–V). Collectively, these *in vitro* findings suggest that HIF-1α loss shifts RORγt⁺ Treg toward a less inflammatory and more suppressive state accompanied by improved mitochondrial integrity, a fusion-associated transcriptional profile, and greater respiratory flexibility. This coordinated functional and metabolic phenotype further suggests that enhanced mitochondrial fitness may help sustain the increased suppressive capacity of ΔHif1a RORγt⁺ Treg.

### HIF-1α deletion limits colitis-associated tumorigenesis and preserves oxidative capacity in colonic Treg

Long-standing colonic IBD increases colorectal cancer incidence and mortality, and cumulative histologic inflammation predicts colorectal neoplasia^69–71^. Because HIF-1α deletion limited both acute and T cell–driven colitis, and inflammatory RORγt⁺ Treg expand during IBD and early dysplastic progression^44^, we asked whether the same mechanism influenced colitis-associated tumorigenesis. WT and ΔHif1a mice received AOM followed by three cycles of 2% DSS and were evaluated through day 63 (Fig. 5A). Serial colonoscopy showed fewer macroscopic lesions in ΔHif1a mice after the first DSS cycle and at endpoint (Fig. 5B). ΔHif1a mice also had longer colons and fewer tumors at necropsy (Fig. 5C,D). In WT CLP, CAC expanded RORγt⁺ Treg; this expansion was markedly blunted by Hif1a deletion without a comparable reduction in total Foxp3⁺ Treg (Fig. 5E). Residual ΔHif1a RORγt⁺ Treg expressed more CTLA-4 and PD-1 and less IL-17A and IFN-γ (Fig. 5F–M), linking the lower tumor burden to reduced accumulation of an inflammatory regulatory state.

**Figure 5.**
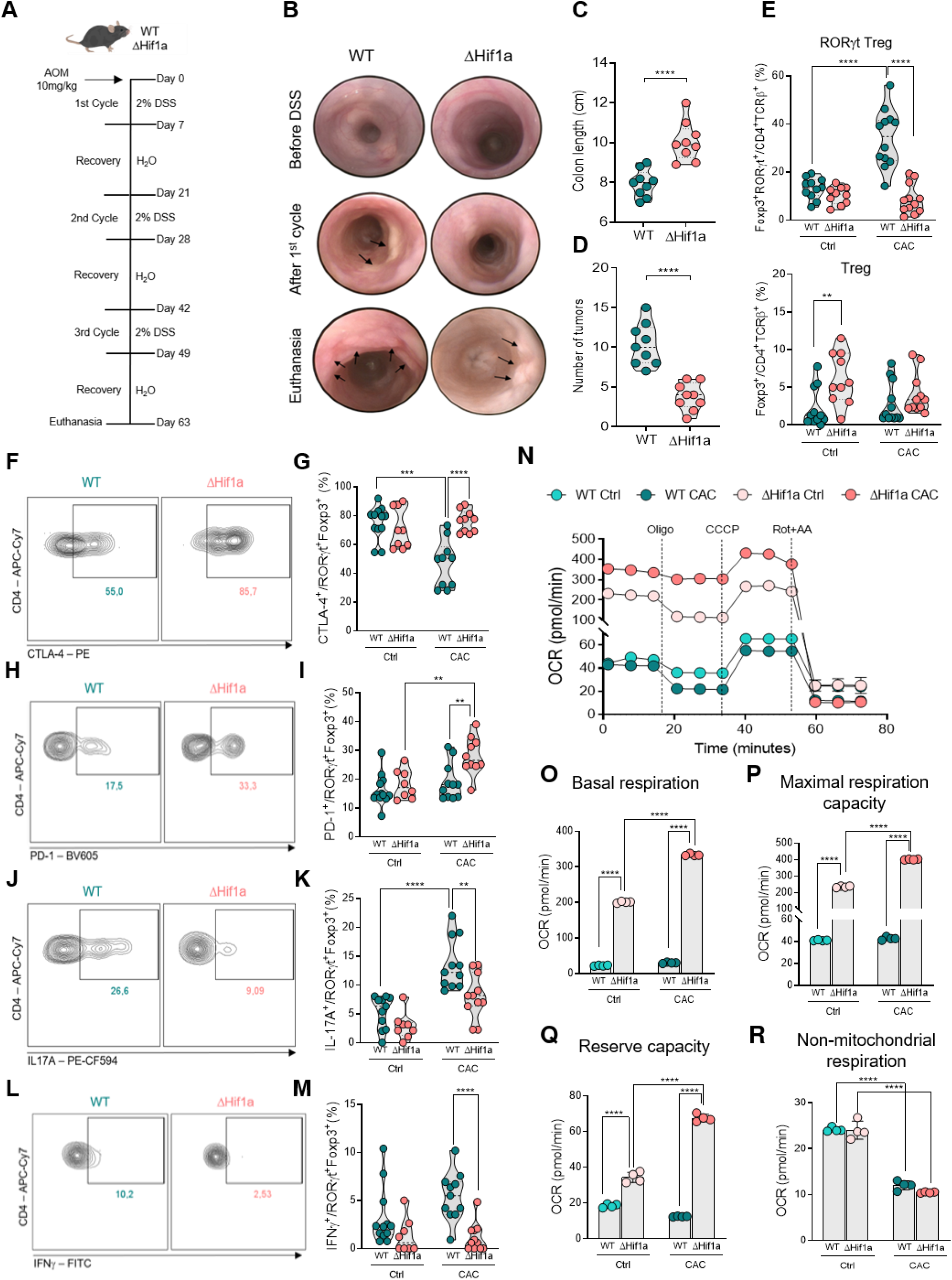
HIF-1α deletion limits colitis-associated tumorigenesis and preserves oxidative capacity in colonic Treg. (A) AOM/DSS CAC protocol with three 7-day cycles of 2% DSS and endpoint on day 63. (B) Representative colonoscopy at baseline, after the first DSS cycle, and at endpoint; arrows indicate lesions. (C) Colon length. (D) Tumor number. (E) Frequencies of CLP Foxp3⁺RORγt⁺ Treg and total Foxp3⁺ Treg under control and CAC conditions. (F–M) Representative plots and quantification of CTLA-4 (F,G), PD-1 (H,I), IL-17A (J,K), and IFN-γ (L,M) in CLP RORγt⁺ Treg. (N) OCR of CLP CCR6⁺ Treg from WT and ΔHif1a mice under control and CAC conditions. (O–R) Basal respiration (O), maximal respiratory capacity (P), reserve capacity (Q), and non-mitochondrial respiration (R). Points represent mice or independently sorted samples; data are mean ± SEM.

Secondary lymphoid organs separated changes in cell abundance from changes in phenotype. In MLN, the frequencies of total Foxp3⁺ Treg, RORγt⁺ Treg, and Th17 cells were broadly comparable across genotype and treatment (Fig. S5A). Nevertheless, ΔHif1a RORγt⁺ Treg expressed more PD-1 and less IL-17A than WT cells during CAC, whereas CTLA-4, IFN-γ, and CD44 did not differ detectably (Fig. S5D). In spleen, WT CAC induced a marked expansion of RORγt⁺ Treg that was blunted by Hif1a deletion without a parallel change in total Foxp3⁺ Treg or Th17 cells (Fig. S5B). The remaining ΔHif1a splenic RORγt⁺ Treg expressed more CTLA-4, PD-1, and CD44 and less IL-17A, while IFN-γ was not significantly altered (Fig. S5E). Thus, HIF-1α loss limited the systemic CAC-associated expansion of this population while preserving a less inflammatory phenotype among the cells that remained.

Homing-receptor analysis showed that this redistribution was selective rather than a generalized loss of CCR9 and CCR6. WT CAC markedly increased the splenic CD103⁺CCR9⁺ fraction of RORγt⁺ Treg, and this expansion was attenuated in ΔHif1a mice; by contrast, splenic CD103⁺CCR6⁺ frequencies remained similar across groups (Fig. S6A, B, D). In CLP, neither the CD103⁺CCR9⁺ nor the CD103⁺CCR6⁺ fraction differed detectably by genotype during CAC (Fig. S6C, E). Because CCR9 preferentially supports small-intestinal homing, whereas the CCR6 axis promotes Treg recruitment and accumulation in the inflamed colon^,73^, the selective splenic result is consistent with reduced expansion or maintenance of a circulating gut-homing reservoir. This interpretation also fits the reported expansion of RORγt⁺ Treg during intestinal inflammation and dysplastic progression^44,62^. In Apc^Min^ mice, which develop adenomas spontaneously, RORγt⁺ Treg did not expand detectably in CLP, MLN, or spleen (Fig. S6F, G), further linking the HIF-dependent redistribution to chronic inflammation rather than neoplasia alone.

We finally asked whether the oxidative phenotype observed *in vitro* persisted in diseased tissue. CCR6⁺ Treg sorted from the CLP of control and CAC mice showed markedly higher oxygen consumption after HIF-1α deletion under both conditions (Fig. 5N). Basal respiration, maximal respiratory capacity, and reserve capacity increased, whereas non-mitochondrial respiration decreased (Fig. 5O–R). The *ex vivo* flux data therefore corroborate the *in vitro* and scATAC-directed findings and associate preserved oxidative fitness with protection from inflammation-driven tumorigenesis.

### Trajectory and regulatory-network analyses position HIF1A along the inflammatory Treg transition

We returned to the human dataset to ask where HIF1A might act along the disease-associated transition. Pseudotemporal ordering resolved seven states across a branched trajectory (Fig. 6A). Mapping cluster identity and condition onto the same structure showed Crohn’s disease cells enriched along branches containing clusters 4–6, whereas healthy cells predominated in other regions (Fig. 6B). Hypoxia and TNFα module scores increased along overlapping portions of the disease-associated branches (Fig. 6C), connecting the trajectory to the pro-inflammatory FOXP3⁺ populations described in the source study^62^.

**Figure 6.**
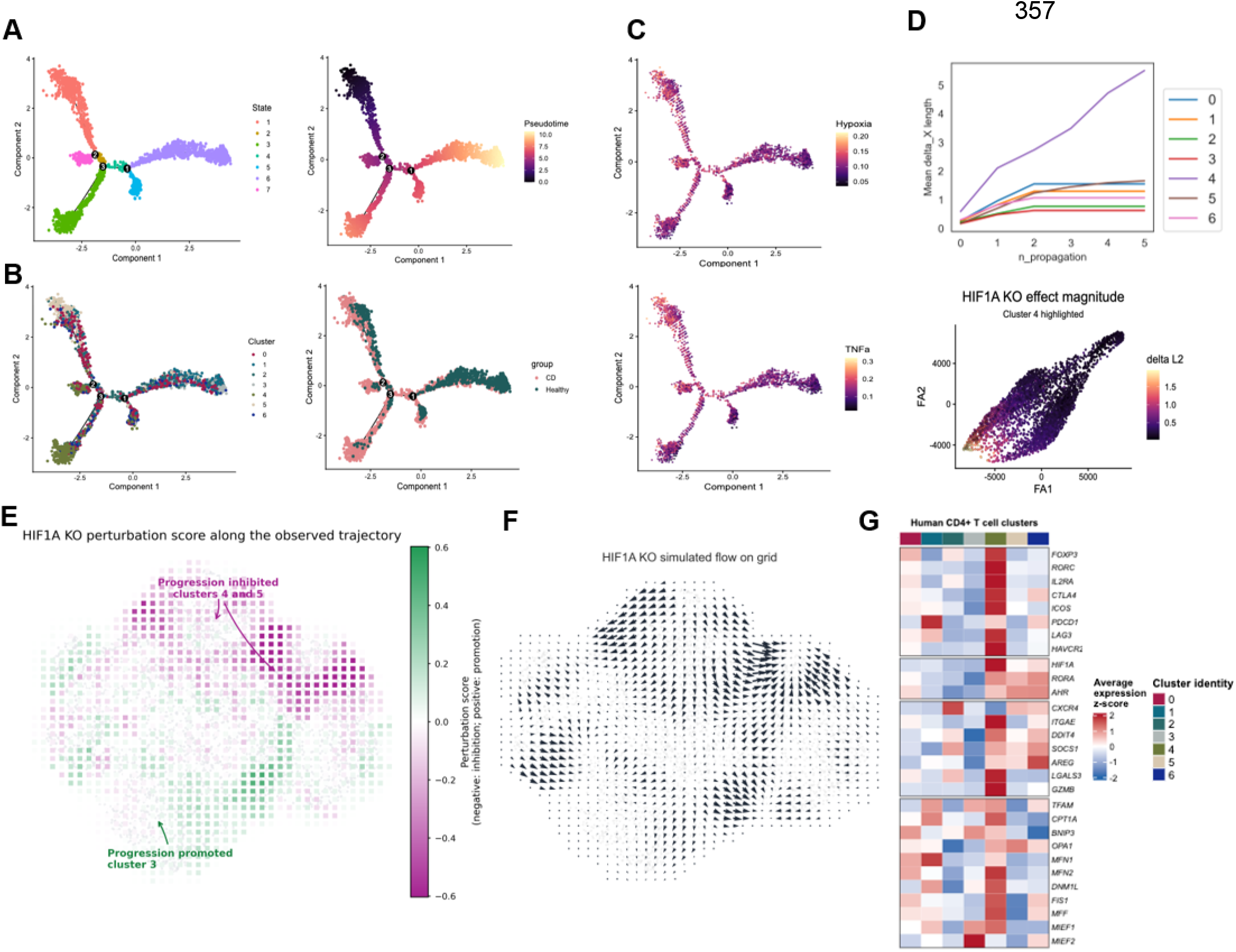
Trajectory and regulatory-network analyses position HIF1A along the inflammatory FOXP3⁺ T cell transition. (A) Branched trajectory colored by inferred state and pseudotime. (B) The same trajectory colored by Seurat cluster and disease group. (C) Hypoxia and TNFα Hallmark module scores projected along the trajectory. (D) Mean propagated HIF1A-knockout expression-shift magnitude by cluster (top) and cell-level effect magnitude with cluster 4 highlighted (bottom). (E) HIF1A-knockout perturbation score along the observed trajectory; positive values align with and negative values oppose the pseudotime gradient. (F) Simulated HIF1A-knockout vector field projected onto the trajectory grid. (G) Average scaled expression of regulatory, inflammatory-adaptation, HIF-associated, and mitochondrial-dynamics genes across clusters. Computational perturbations are predictions rather than measurements in HIF1A-deficient human cells.

We next inferred cluster-specific gene-regulatory networks with CellOracle and simulated HIF1A loss^64^. Cluster 4 showed the largest mean propagated expression-shift magnitude, with cell-level effects also evident across other disease-associated states (Fig. 6D). Comparing simulated HIF1A-knockout vectors with the observed pseudotime gradient yielded negative perturbation scores in clusters 4 and 5, consistent with opposition to progression along those branches, and positive scores in cluster 3 (Fig. 6E). Projection of the simulated vector field illustrated state-dependent redirection rather than a uniform collapse of the FOXP3⁺ compartment (Fig. 6F). Because the networks were inferred from observational data, these results are predictions that require perturbation in patient-derived cells.

A cluster-level heatmap placed regulatory, inflammatory-adaptation, HIF-associated, and mitochondrial-dynamics genes within the same transcriptional landscape (Fig. 6G). The convergence of these programs in cluster 4, together with the negative perturbation score predicted for disease-associated branches, mirrors the regulatory and mitochondrial coupling observed after HIF-1α deletion in mice. The cross-species comparison therefore positions HIF1A within selected inflammatory Treg trajectories without implying that every human FOXP3⁺ state will respond identically.

## DISCUSSION

Emerging evidence indicates that intestinal RORγt⁺ Treg are not a fixed regulatory population but can adopt distinct suppressive or inflammatory states according to the surrounding tissue environment^6–27,44^. HIF-1α is a central sensor of oxygen and metabolic stress and is known to influence the balance between Th17 and conventional Treg differentiation^38–41^. However, its specific function in differentiated intestinal RORγt⁺ Treg and its contribution to chronic intestinal inflammation and tumorigenesis have remained unclear. Here, we demonstrate that HIF-1α acts as a context-dependent checkpoint that links inflammatory adaptation to the functional and mitochondrial state of RORγt⁺ Treg. HIF-1α deletion reinforced suppressive activity and oxidative fitness, limited inflammatory cytokine production, and reduced disease severity across acute DSS colitis, T cell transfer colitis, and colitis-associated colorectal cancer (CAC).

The human single-cell analyses provide a translational foundation for the mouse experiments. Kosinsky and colleagues identified Crohn’s disease-associated FOXP3⁺ T cell populations characterized by TNF responsiveness, inflammatory transcription, and impaired suppressive activity^62^. In our reanalysis, HIF1A detection and hypoxia-responsive activity were concentrated within related disease-enriched RORC⁺FOXP3⁺ states rather than being uniformly distributed throughout the regulatory compartment. Hypoxia, TNF-NF-κB, glycolytic, and inflammatory programs converged along selected Crohn’s disease-associated branches, supporting a model in which inflammatory regulatory states emerge through the coordinated action of cytokine and metabolic signals rather than through uniform loss of FOXP3 identity. Importantly, network-based HIF1A perturbation was predicted to oppose progression along specific disease-associated branches^64^. The direction of this predicted response resembled the less inflammatory, more regulatory, and metabolically fitter phenotype measured after genetic HIF-1α deletion in mice, providing cross-species support for a conserved HIF-associated program. Nevertheless, these human perturbations remain computational predictions and require direct validation in patient-derived cells.

The effects of HIF-1α on Treg have differed across experimental systems, emphasizing the importance of cellular and environmental context. HIF-1α can bind regulatory regions of Foxp3 and promote Treg accumulation and function during mucosal hypoxia; accordingly, HIF-1α-deficient CD4⁺CD25⁺ cells were previously reported to provide impaired protection in transfer colitis^76^. Conversely, persistent HIF-1α stabilization caused by Foxp3-restricted Vhl deletion destabilized the regulatory phenotype, increased IFN-γ production, and impaired suppression, effects that were reversed by additional HIF-1α deletion^77^. Sustained systemic hypoxia has likewise been associated with reduced regulatory features and emergence of exTreg-Th17 cells^63^. Our findings refine this apparent paradox by focusing on cells with a history of RORγt expression. In this compartment, HIF-1α was dispensable for normal lymphocyte development under homeostatic conditions but favored inflammatory and metabolically stressed states once intestinal inflammation was established.

During acute DSS colitis, protection in ΔHif1a mice was not explained by elimination of RORγt⁺ Treg. Instead, HIF-1α deletion altered their phenotype: ΔHif1a RORγt⁺ Treg expressed more CTLA-4, produced less IFN-γ and IL-17A, and were associated with increased IL-10 in diseased colonic tissue. In the spleen, ΔHif1a RORγt⁺ Treg accumulated during colitis without a corresponding increase in inflammatory cytokine production, indicating that HIF-1α loss uncoupled extraintestinal accumulation from acquisition of an inflammatory phenotype. Broad profiling of ILC3, γδ T cell, dendritic cell, monocyte, neutrophil, and macrophage compartments did not reveal a compositional change proportional to the degree of protection. Although this finding does not exclude functional changes in other RORγt-lineage populations, it directs attention to altered regulatory quality rather than a generalized reduction in inflammatory-cell abundance.

Consistent with the *in vivo* experiments, ΔHif1a RORγt⁺ Treg displayed enhanced regulatory function *in vitro*. These cells secreted more IL-10 and less IL-17A, expressed higher levels of inhibitory receptors, and more efficiently restrained responder T cell proliferation across multiple Treg:responder ratios. The adoptive-transfer model provided complementary evidence *in vivo*. Rag1^⁻/⁻^ recipients lacked endogenous T and B cells and received the same WT pathogenic naïve CD4⁺ T cell inoculum, while the genotype of the cotransferred Treg population constituted the principal experimental variable. Under these standardized conditions, ΔHif1a Treg more effectively preserved body weight and colon length, limited splenomegaly and tissue injury, and reduced inflammatory effector T cell responses. The recipient myeloid compartment was not removed and may have participated in the protection; however, it was held constant across cotransfer groups. Together, the suppression and transfer experiments support an enhanced regulatory capacity of ΔHif1a Treg rather than protection arising solely from broader developmental effects of Rorc^Cre^-mediated deletion.

Metabolic programming is an active determinant of Treg function. Foxp3 suppresses Myc-driven glycolysis and promotes oxidative metabolism, enabling Treg to function in low-glucose, high-lactate environments^79^, while mitochondrial complex III is required to maintain suppressive gene expression even when Treg abundance and Foxp3 expression remain intact^80^. In our study, mouse colonic Treg scATAC-seq showed mitochondrial-maintenance and oxidative programs. Genetic HIF-1α deletion subsequently reduced the frequency of cells with dysfunctional mitochondria and mitochondrial ROS, increased MitoTracker Red signal per cell, enhanced expression of Tfam, Mfn1, and Mfn2, and reduced Drp1, Fis1, and Fis2. These changes were accompanied by greater basal and maximal respiration and reserve capacity both *in vitro* and in Treg isolated from diseased tissue. The findings extend our previous demonstration that *in vitro*-generated RORγt⁺ Treg are highly suppressive and dependent on oxidative phosphorylation^74^. Although the coordinated increase in mitochondrial integrity, fusion-associated transcription, and respiratory flexibility provides a plausible energetic basis for enhanced suppression, direct perturbation of mitochondrial dynamics will be required to determine whether these changes are necessary or sufficient for the regulatory phenotype.

The protection observed in CAC expands the relevance of this mechanism beyond acute inflammation. Patients with long-standing colonic IBD have an increased risk of colorectal cancer, and cumulative histological inflammation is an independent predictor of neoplasia^69–71^. In the AOM/DSS model, HIF-1α deletion reduced macroscopic tumor burden and restricted the inflammation-associated expansion of RORγt⁺ Treg while preserving a less inflammatory and more oxidative phenotype among the remaining cells. This is consistent with evidence that RORγt⁺ or IL-17-producing Treg can accumulate and acquire tumor-promoting properties during intestinal inflammation and dysplastic progression^12,16,27,43,44^. The absence of comparable RORγt⁺ Treg expansion in Apc^Min^ mice further indicates that repeated mucosal inflammation, rather than neoplasia alone, is a major driver of the HIF-dependent state.

Collectively, these findings support a model in which inflammatory cytokines, tissue hypoxia, and metabolic stress converge on HIF-1α in intestinal RORγt⁺ Treg during persistent inflammation (Fig. 7). Sustained HIF-associated activity favors an inflammatory regulatory state characterized by IL-17A and IFN-γ production, mitochondrial dysfunction and ROS accumulation, fission-associated transcription, and limited oxidative reserve, together with inflammation-dependent redistribution and expansion of the population. HIF-1α deletion shifts this balance toward mitochondrial maintenance and fusion, greater respiratory flexibility, increased IL-10 and inhibitory-receptor expression, and stronger suppression. This coordinated change limits effector T cell activation, intestinal tissue injury, and inflammation-associated tumorigenesis. The predicted redirection of HIF1A-perturbed RORC⁺FOXP3⁺ cells from patients with Crohn’s disease toward a similar regulatory and metabolic state provides the human counterpart of the proposed model.

**Figure 7.**
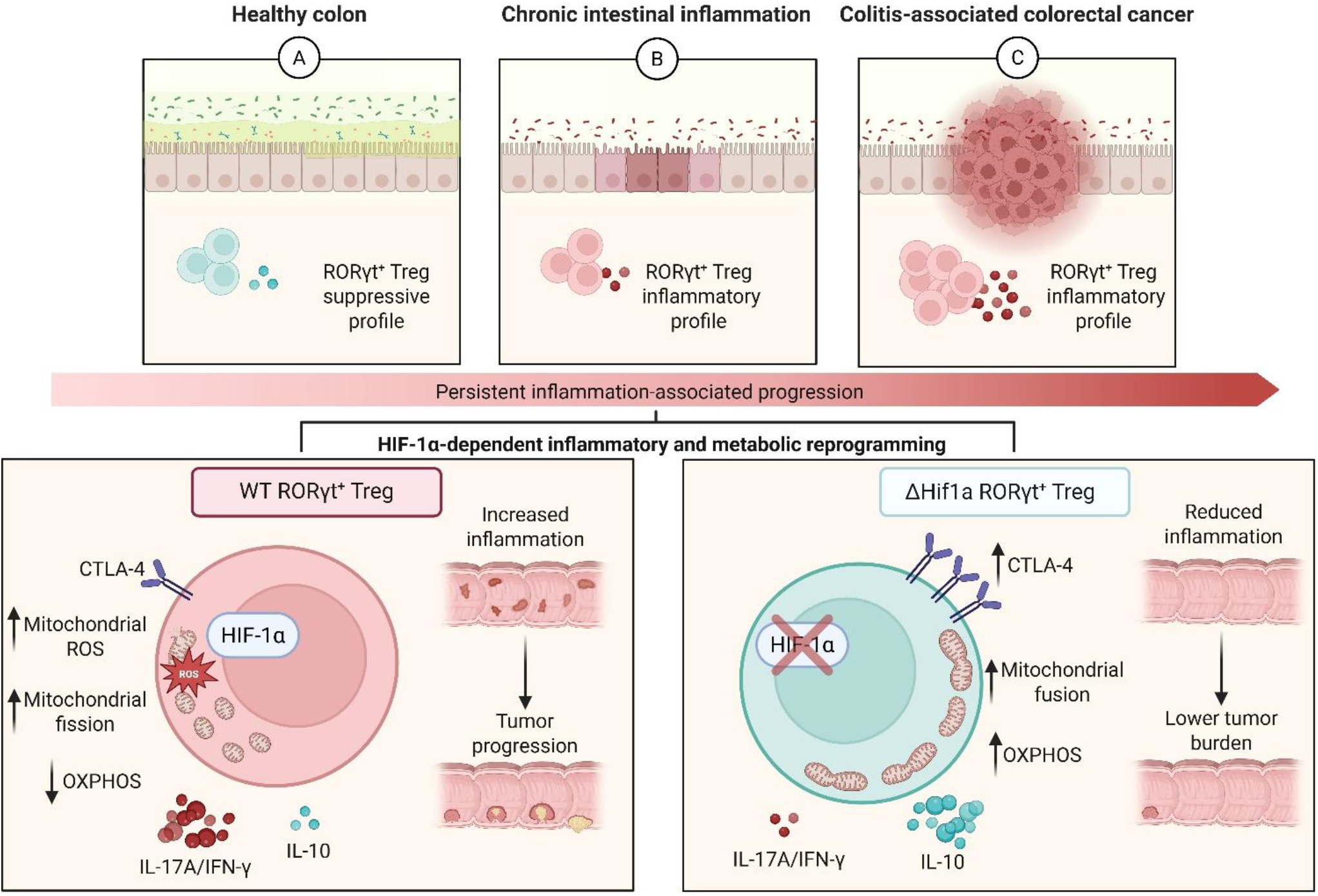
Proposed model of HIF-1α-dependent inflammatory and metabolic reprogramming of RORγt⁺ regulatory T cells during chronic intestinal inflammation and colitis-associated colorectal cancer. (A) Under intestinal homeostasis, RORγt⁺ regulatory T (Treg) cells maintain a predominantly suppressive phenotype and contribute to mucosal immune tolerance and epithelial barrier integrity. (B) During chronic intestinal inflammation, persistent environmental and inflammatory signals promote HIF-1α-dependent functional and metabolic reprogramming of RORγt⁺ Treg cells toward a less suppressive and more inflammatory state. This phenotype is characterized by increased mitochondrial reactive oxygen species (ROS) and mitochondrial fission, reduced oxidative phosphorylation (OXPHOS), and enhanced production of the inflammatory cytokines IL-17A and IFN-γ, accompanied by reduced IL-10 production and CTLA-4 expression. (C) Persistence of this inflammatory program may contribute to epithelial damage and support the development and progression of colitis-associated colorectal cancer. By contrast, HIF-1α deletion in RORγt⁺ cells preserve a regulatory and metabolically oxidative phenotype, characterized by enhanced mitochondrial fusion and OXPHOS, increased CTLA-4 and IL-10 expression, and reduced IL-17A and IFN-γ production. These changes restrain intestinal inflammation and are associated with reduced tumor burden. The illustration summarizes the proposed context-dependent role of HIF-1α in controlling the functional plasticity and mitochondrial fitness of intestinal RORγt⁺ Treg cells. ROS, reactive oxygen species; OXPHOS, oxidative phosphorylation; CAC, colitis-associated colorectal cancer.

The similarity between the predicted response of patient-derived FOXP3⁺ cells and the experimentally observed ΔHif1a phenotype in mice highlights a potential path toward application. Patient-derived CD45RA⁺ Treg can be expanded *ex vivo* while retaining suppressive activity against lymphocytes from inflamed Crohn’s disease tissue^81^, and administration of antigen-specific autologous Treg has been shown to be feasible and generally well tolerated in patients with refractory Crohn’s disease^82^. These observations raise the possibility that transient HIF-1α modulation during *ex vivo* Treg expansion or engineering could improve mitochondrial fitness and suppressive stability before adoptive transfer. The HIF1A-associated transcriptional signature may also help identify patients or regulatory-cell states most likely to benefit from such an approach. This strategy will require direct testing in patient-derived RORC⁺ Treg, including measurements of cytokine production, suppressive function, lineage stability, and mitochondrial respiration after HIF1A modulation.

In conclusion, we demonstrate that HIF-1α connects hypoxia-responsive transcription to the inflammatory, suppressive, and mitochondrial states of intestinal RORγt⁺ Treg. Limiting this program enhanced regulatory function and oxidative reserve and reduced both intestinal inflammation and inflammation-associated tumorigenesis. Because epithelial HIF-1α supports barrier protection during experimental colitis^78^ and HIF signaling has essential functions across multiple tissues, systemic HIF-1α blockade would be unlikely to provide the required specificity. The cross-species convergence observed here instead supports the development of cell-state-restricted or *ex vivo* strategies aimed at reprogramming dysfunctional RORγt⁺ Treg. Demonstrating that selective HIF1A modulation reproduces the murine suppressive and mitochondrial phenotype in cells from patients with IBD represents the critical next step toward therapeutic translation.

## MATERIALS AND METHODS

### Study design

This study tested how HIF-1α regulates the phenotype, suppressive function, and metabolism of RORγt⁺ Treg during intestinal inflammation. The experimental sequence combined human scRNA-seq reanalysis, conditional genetic deletion, acute DSS colitis, AOM/DSS-induced CAC, and T cell transfer colitis. Mouse colonic Treg scATAC-seq was then used to nominate cell-intrinsic regulatory and mitochondrial outputs, which were tested by *in vitro* differentiation, suppression, cytokine assays, mitochondrial dyes, RT–qPCR, and extracellular-flux analysis. Human trajectory inference and gene-regulatory-network perturbation provided a final cross-species test of the model. Sample sizes were guided by prior studies and pilot experiments; no formal power calculation was performed. Mice were assigned according to littermate availability, and histologic scoring and data analysis were performed blinded when feasible. Points in the figures denote individual animals, independent cultures, or independently sorted samples as specified.

### Animals

Male mice 8–10 weeks of age and 18–25 g were used unless indicated otherwise. Hif1a^flox/flox^ mice were crossed with Rorc^Cre^ mice (The Jackson Laboratory) to generate Rorc^Cre^ Hif1a^flox/flox^ mice (ΔHif1a); Rorc^Cre^ littermates served as WT controls. Apc^Min^, Rag1^⁻/⁻^, and C57BL/6 mice were used in the indicated experiments. Animals were maintained under specific pathogen–free conditions on a 12-hour light/dark cycle with sterile food and water ad libitum. Procedures complied with Brazilian Federal Law 11,794/08 and were approved by the Ethics Committee on Animal Use of the University of São Paulo (protocol 7288101218).

### Acute DSS colitis and colitis-associated colorectal cancer

For acute colitis, WT and ΔHif1a mice received 2% (w/v) DSS in drinking water from day 0 through day 7, followed by regular water until euthanasia on day 10. Body weight, stool consistency, visible bleeding, and behavior were monitored daily and combined using the clinical score in table S1.

For CAC, mice received azoxymethane (AOM; 10 mg/kg, intraperitoneally). After 48 hours, 2% DSS was provided for seven days in three cycles (days 0–7, 21–28, and 42–49 relative to the first DSS exposure), with water during recovery intervals. Animals were monitored through day 63. Colon length and macroscopic tumors were recorded at necropsy. High-resolution colonoscopy was performed under anesthesia before DSS, after the first DSS cycle, and at endpoint to document lesion development.

### Adoptive transfer colitis

CD4⁺ T cell subsets were isolated from pooled spleens and lymph nodes. Treg were sorted as CD4⁺CD25^high^ cells from WT or ΔHif1a donors. Naïve CD4⁺ T cells (CD4⁺CD45RB^high^CD8⁻CD11c⁻CD19⁻) were sorted from WT mice. Each Rag1⁻/⁻ recipient received 2.8 × 10^5^ naïve cells alone or together with 7 × 10^4^ WT or ΔHif1a Treg (4:1 ratio) by intraperitoneal injection in sterile PBS. PBS-only controls were included. Body weight and clinical condition were monitored for five weeks before tissue collection.

### Histology

Large-intestinal segments were fixed for 24 hours at 4°C in Methacarn (60% methanol, 30% chloroform, and 10% glacial acetic acid), transferred to 70% ethanol, paraffin embedded, sectioned, and stained with hematoxylin and eosin. Images were acquired at 10×, 20×, and 40× on a Nikon microscope. Colonic inflammation and tissue injury were scored using established histomorphologic criteria^61^.

### Cell isolation and flow cytometry

Thymus, spleen, MLN, PLN, and colon were collected into ice-cold PBS with 2% fetal bovine serum (FBS). Spleens and lymph nodes were mechanically dissociated through 70-µm strainers; erythrocytes were lysed for 3 minutes before filtration through 40-µm strainers. Colons were opened, washed, and cut into approximately 1-cm fragments. Epithelial cells were removed by incubation in RPMI containing 3% FBS, 5 mM EDTA, and 0.145 mg/ml dithiothreitol for 20 minutes at 37°C with agitation. Remaining tissue was digested in RPMI containing 0.5 mg/ml collagenase VIII and 10 U/ml DNase for 20 minutes at 37°C, filtered, washed, and resuspended for staining.

Viability, surface, cytokine, and transcription-factor staining followed standard manufacturer protocols. Treg, RORγt⁺ Treg, and Th17 cells were defined within live CD45⁺TCRβ⁺CD4⁺ cells by Foxp3 and RORγt. Intracellular cytokines were assessed after stimulation in the presence of secretion inhibitors. Innate and innate-like populations were defined using the sequential gates in Fig. S3. Antibodies, fluorochromes, clones, manufacturers, and working dilutions are listed in tables S2–S4. Samples were acquired on BD FACSCanto or FACSAria instruments with FACSDiva and analyzed in FlowJo.

### *In vitro* differentiation of Treg and RORγt⁺ Treg

Naïve CD4⁺ T cells were isolated from spleen and lymph nodes by sorting CD4⁺CD44^low– int^CD62L⁺ cells or with the EasySep Mouse Naïve CD4⁺ T Cell Isolation Kit. Cells (2 × 10^5^ per well) were stimulated with plate-bound anti-CD3 (2 µg/ml) and soluble anti-CD28 (1 µg/ml) in complete RPMI. Treg polarization used TGF-β (5 ng/ml), IL-2 (50 U/ml), anti-IL-12/23p40 (1 µg/ml), anti-IL-4 (1 µg/ml), and anti-IFN-γ (1 µg/ml) for four days. For RORγt⁺ Treg differentiation, cultures were transferred to freshly coated plates on day 4 and restimulated with anti-CD3 and anti-CD28 plus IL-6 (2.5 ng/ml) and TGF-β (5 ng/ml) for four additional days. Reagents are detailed in table S5. Foxp3 and RORγt were measured by flow cytometry at endpoint.

### Treg suppression assay

The assay was adapted from the CellTrace Violet suppression protocol of Ellestad and Anderson^68^ and the *in vitro* RORγt⁺ Treg workflow described previously^74^. T CD4+ responder cells were labeled with CellTrace Violet, and 7.5 × 10⁴ cells were plated per well in U-bottom 96-well plates. *In vitro*–generated WT or ΔHif1a Foxp3⁺RORγt⁺ Treg were added at Treg:responder ratios of 0:1, 1:16, 1:8, 1:4, 1:2, and 1:1, togheter with 25.000 splenocytes from RAG^-/-^ mice. Cultures were stimulated with soluble anti-CD3 (1 µg/ml) for 72 hours at 37°C and 5% CO₂. Cells were then stained for viability and responder proliferation was quantified as CellTrace Violet dilution by flow cytometry. The fraction of responders that completed at least one division was expressed relative to stimulated responder splenocytes cultured without Treg.

### ELISA

IL-10 and IL-17A were quantified in culture supernatants or tissue lysates with R&D Systems kits according to the manufacturer’s instructions. Concentrations were calculated from standard curves generated on the same plate.

### Mitochondrial mass, membrane potential, and reactive oxygen species

Mitochondrial mass and membrane potential were assessed by combined MitoTracker Green and MitoTracker Red staining. Cells with high MitoTracker Green and low MitoTracker Red signal were quantified as the dysfunctional-mitochondria gate shown in Fig. 4G. Mitochondrial reactive oxygen species were measured with MitoSOX Red. Staining was performed in live cells according to manufacturer instructions, followed by immediate flow-cytometric acquisition with matched unstained controls.

### Confocal microscopy and mask-based image analysis

*In vitro*–differentiated WT and ΔHif1a RORγt⁺ Treg were stained with SYBR Green to identify nuclei and MitoTracker Red to visualize mitochondria, according to the manufacturers’ instructions, and immediately imaged by confocal microscopy. Transmitted-light, SYBR Green, and MitoTracker Red channels were acquired separately using the same magnification and acquisition settings for all cultures. Channel-specific images were retained for quantitative analysis and merged for visualization. Quantification was performed using a fixed sequential mask-based workflow (Fig. S5A–D). First, cellular objects were segmented in transmitted-light images (Fig. S5A), and individual objects were assigned pseudocolored masks (Fig. S5B). Fluorescence images were subsequently used to identify SYBR Green–positive nuclei and define the corresponding cell-associated regions retained for analysis (Fig. S5C). A final per-cell region of interest was generated for each selected object and overlaid on the merged SYBR Green and MitoTracker Red channels (Fig. S5D). Debris, cell aggregates, poorly segmented objects, and objects intersecting the image borders were excluded using fixed size and morphology criteria. Identical intensity thresholds, segmentation parameters, and exclusion criteria were applied to WT and ΔHif1a images. The number of selected nuclei was recorded as a quality-control measure of cell representation, whereas the mitochondrial signal was calculated as the mean MitoTracker Red fluorescence intensity within each selected cell. Measurements from the acquired fields were averaged to obtain one value for each independent culture.

### Extracellular flux analysis

Oxygen consumption rate (OCR) was measured with an XF96 Extracellular Flux Analyzer (Agilent). *In vitro*–differentiated Treg or RORγt⁺ Treg, or CCR6⁺ Treg sorted from CLP, were plated at 4 × 10^5^ cells per well on XF96 plates coated for 2 hours with poly-D-lysine (100 µg/ml). Assay medium was unbuffered RPMI 1640 supplemented with 2 mM glutamine, 25 mM glucose, and 1 mM sodium pyruvate. Oligomycin (1 µg/ml), CCCP (5 µM), and rotenone (1 µg/ml) plus antimycin A (1 µM) were sequentially injected for the mitochondrial stress test. Basal respiration, maximal respiration, reserve capacity, and non-mitochondrial respiration were calculated and normalized to cell number.

### RT–qPCR

RNA was purified from *in vitro*–differentiated RORγt⁺ Treg with the RNeasy Mini Kit (QIAGEN, 74104). One microgram of RNA was reverse-transcribed with M-MLV reverse transcriptase and oligo(dT) primers. Reactions contained 500 nM of each primer and SYBR Green PCR Master Mix in a 10-µl volume and were run in technical triplicate. Relative expression was calculated by the 2^−ΔΔCt^ method using β-actin as the reference. Primer sequences supplied for *Drp1*, *Opa1*, *Mfn1*, *Mfn2*, *Fis1*, *Fis2*, *Foxp3*, *Rorc*, and *Hif1a* are listed in table S6; *Tfam* was measured using the same workflow.

### Human scRNA-seq reanalysis

Public scRNA-seq data from ileal CD4⁺ T cells of healthy donors and patients with Crohn’s disease were obtained from GEO (GSE209832)^62^. The source study generated count matrices with Cell Ranger 6.1.1. Genes detected in fewer than three cells and cells with fewer than 200 detected genes or greater than 20% mitochondrial transcripts were excluded, consistent with the source analysis. Samples were normalized independently with SCTransform, 3,000 variable genes were used for anchor-based integration, and principal-component and shared-nearest-neighbor analyses were performed in Seurat. Cells in the FOXP3⁺ regulatory compartment were reclustered and visualized with UMAP, yielding clusters 0–6. Cluster and condition proportions were calculated from patient-level cell counts. HIF1A-positive cells were defined by detectable *HIF1A* transcript, and the RORγt-high density map used cells with detectable *RORC* together with a positive RORγt-associated module score.

For sample-aware pathway analysis, normalized expression was aggregated by patient and cluster. Hallmark gene sets were obtained from MSigDB. ssGSEA scores were calculated for each patient–cluster combination, standardized as z scores by pathway, and displayed by hierarchical clustering^65^. Differential expression between Crohn’s disease and healthy samples was estimated using patient-level pseudobulk counts and DESeq2 statistics. Preranked GSEA used the DESeq2 ranking statistic and the HALLMARK_HYPOXIA gene set^66^.

### Trajectory inference and *in silico* HIF1A perturbation

Pseudotemporal ordering of the human FOXP3⁺ compartment was performed with Monocle using DDRTree reduction in two dimensions^67^. The root was assigned to the healthy-enriched branch before condition comparison. Cell states, pseudotime, Seurat cluster, and disease group were projected onto the same trajectory. Hypoxia and TNFα scores were calculated from the corresponding Hallmark gene sets and displayed as continuous values along the embedding.

Cluster-specific gene-regulatory networks were inferred with CellOracle using the human promoter base regulatory network and normalized scRNA-seq expression^64^. For the *in silico* knockout, *HIF1A* expression was set to zero and the predicted expression shift was propagated iteratively through each cluster-specific network. The Euclidean length of the simulated expression-shift vector was summarized across propagation steps and projected into the trajectory embedding as a vector field. A pseudotime-gradient vector field represented observed progression. The perturbation score was calculated as the local inner product between the simulated HIF1A-knockout vector and the pseudotime-gradient vector; positive values indicate alignment with observed progression and negative values indicate opposition. These simulations predict changes in cell identity and do not estimate post-perturbation cell numbers.

### Mouse single-cell ATAC-seq reanalysis

Mouse colonic Treg scATAC-seq data were obtained from GEO (GSM6697676)^46^. The peak matrix, barcodes, peaks, fragments, and metadata were analyzed in R with Seurat and Signac. Peaks were converted to GRanges using the mm10 genome annotation. Quality control included fragment depth, nucleosome signal, transcription-start-site enrichment, fraction of reads in peaks, and removal of low-complexity or atypically fragmented libraries. Colonic Treg were normalized by TF–IDF, reduced by latent semantic indexing, embedded by UMAP, and clustered with a shared-nearest-neighbor graph.

Gene activity was inferred with Signac GeneActivity and used to classify RORγt⁺ and RORγt⁻ states. Coverage plots compared accessibility at selected loci. JASPAR2020 motifs were mapped to peaks, and chromVAR was used to calculate motif-deviation scores. Within RORγt⁺ Treg, HIF1A motif-high and motif-low groups were defined from the selected HIF1A motif-deviation score; these labels indicate relative motif-associated accessibility and not HIF-1α protein abundance or expression. Differential accessibility was tested by logistic regression controlling for ATAC depth, and peaks were annotated to nearby genes to visualize suppressive, phenotypic, and metabolic programs.

## Statistical analysis

Experimental data are shown as mean ± SEM unless otherwise indicated. Two-group comparisons used two-tailed unpaired Student’s t tests. Comparisons among three or more groups used one-or two-way ANOVA with multiplicity-adjusted post hoc tests as appropriate; longitudinal measurements were analyzed by two-way repeated-measures ANOVA. Tests were performed in GraphPad Prism. Exact P values are shown when available; otherwise, *P < 0.05, **P < 0.01, ***P < 0.001, and ****P < 0.0001. Individual points denote biological replicates. No statistical test was used to infer causality from the human observational or *in silico* analyses.

## Acknowledgments

We thank the CEFAP-USP for its support with cell sorting for the *in vitro* experiments.

## Data and materials availability

The human scRNA-seq and mouse scATAC-seq source data are available through GEO under accessions GSE209832 and GSM6697676, respectively. All other data are available in the main text or supplementary materials.

## SUPPLEMENTARY FIGURES

**Figure S1.**
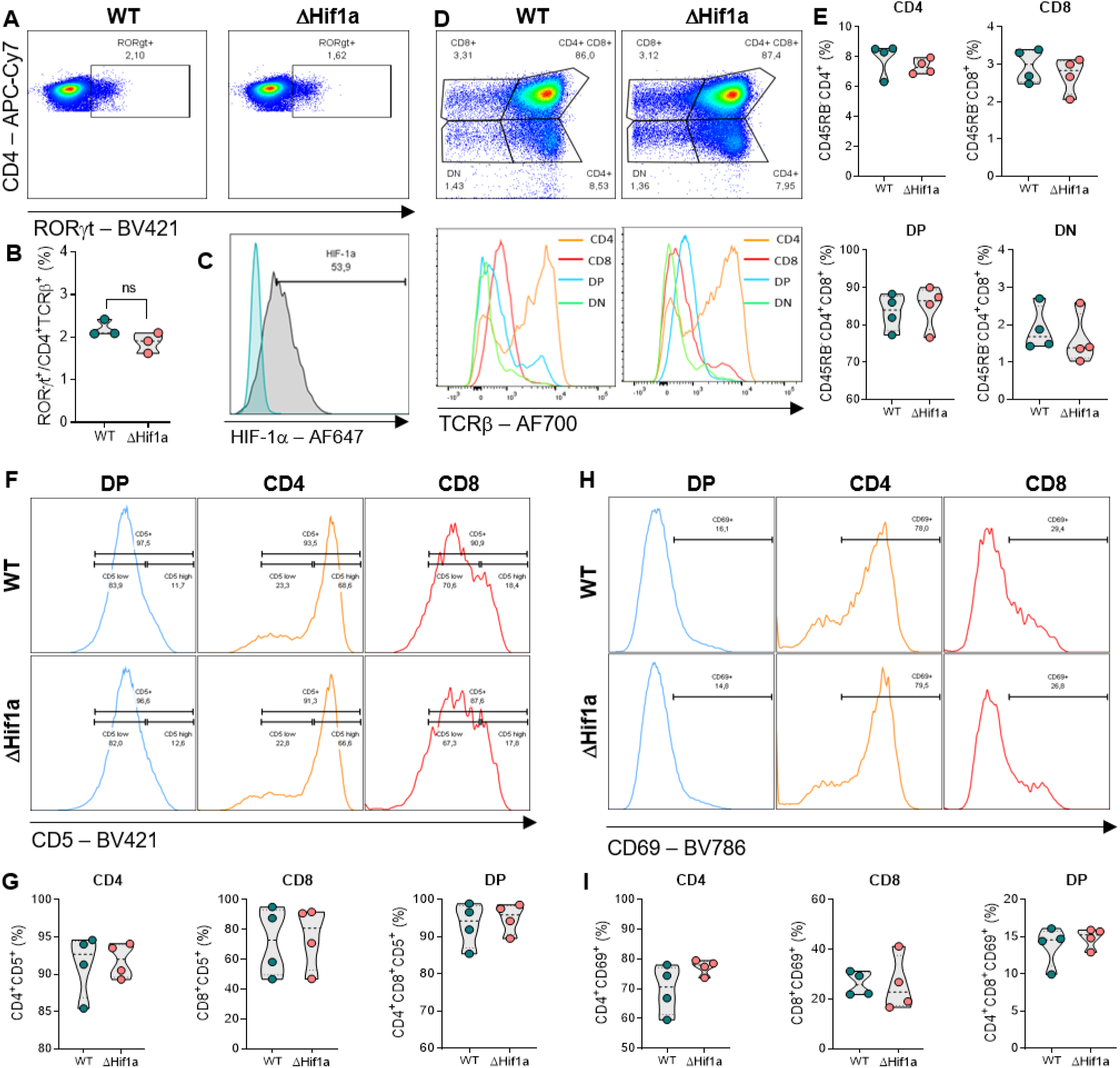
Thymic development and peripheral homeostasis are broadly preserved in ΔHif1a mice. (A,B) Representative flow cytometry and frequencies of RORγt⁺ CD4⁺ T cells in WT and ΔHif1a mice. (C) HIF-1α staining. (D,E) Representative thymic CD4/CD8 subsets and quantification. (F,G) CD5 expression in double-positive, CD4 single-positive, and CD8 single-positive thymocytes. (H,I) CD69 expression in the indicated thymic populations. Points represent mice; data are mean ± SEM.

**Figure S2.**
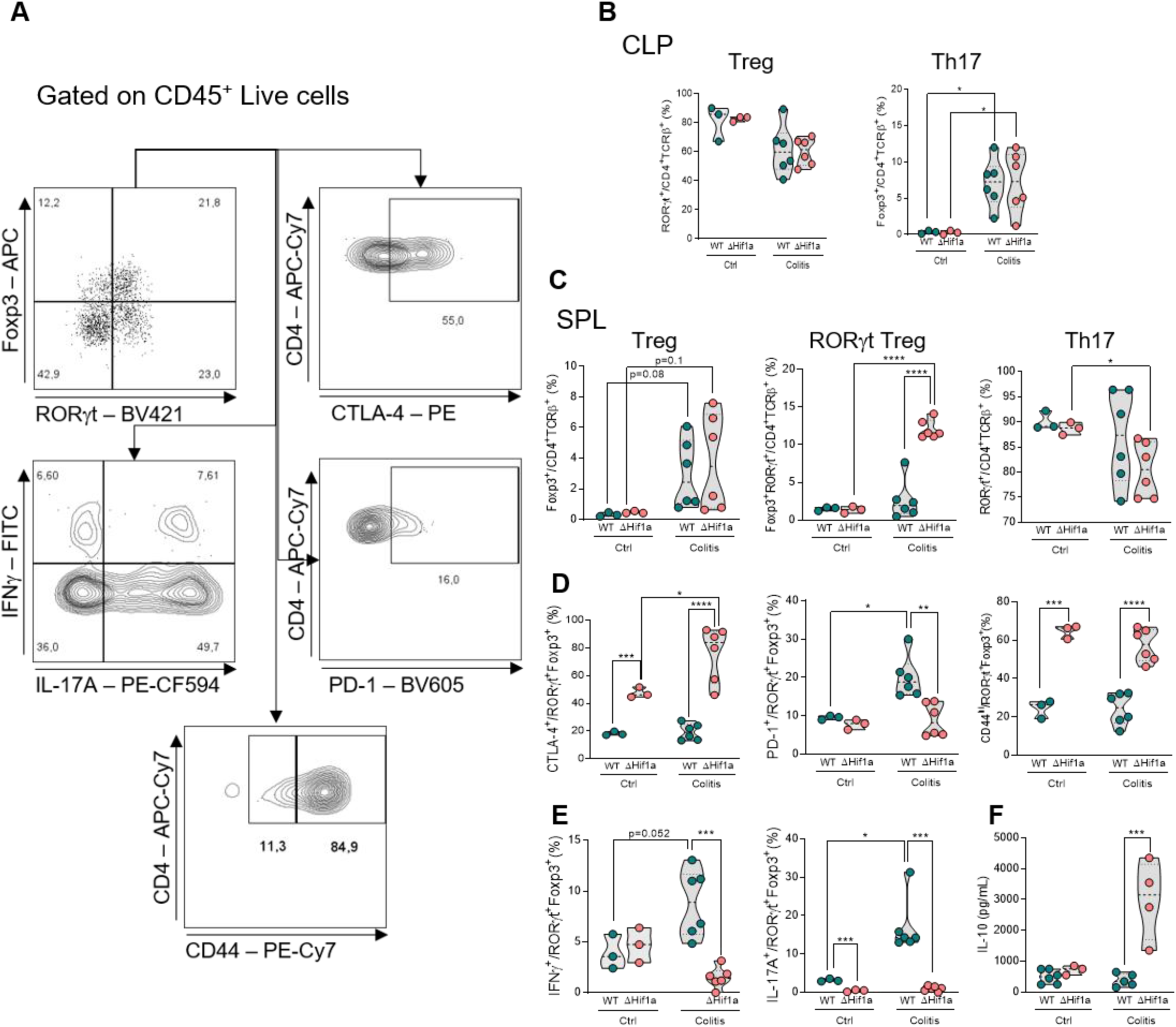
Local and systemic T cell phenotypes and colon IL-10 during acute DSS colitis. (A) Sequential gating of live CD45⁺ CLP cells to identify CD4⁺ T cells, Foxp3⁺ Treg, RORγt⁺ Treg, Th17 cells, and CTLA-4, PD-1, CD44, IFN-γ, and IL-17A. (B) Frequencies of CLP Foxp3⁺ Treg and Th17 cells. (C) Frequencies of Foxp3⁺ Treg, Foxp3⁺RORγt⁺ Treg, and Th17 cells in spleen. (D) CTLA-4, PD-1, and CD44 in splenic RORγt⁺ Treg. (E) IFN-γ and IL-17A in splenic RORγt⁺ Treg. (F) IL-10 concentration in colon tissue. Groups are WT and ΔHif1a mice under control or DSS-colitis conditions. Points represent mice; data are mean ± SEM.

**Figure S3.**
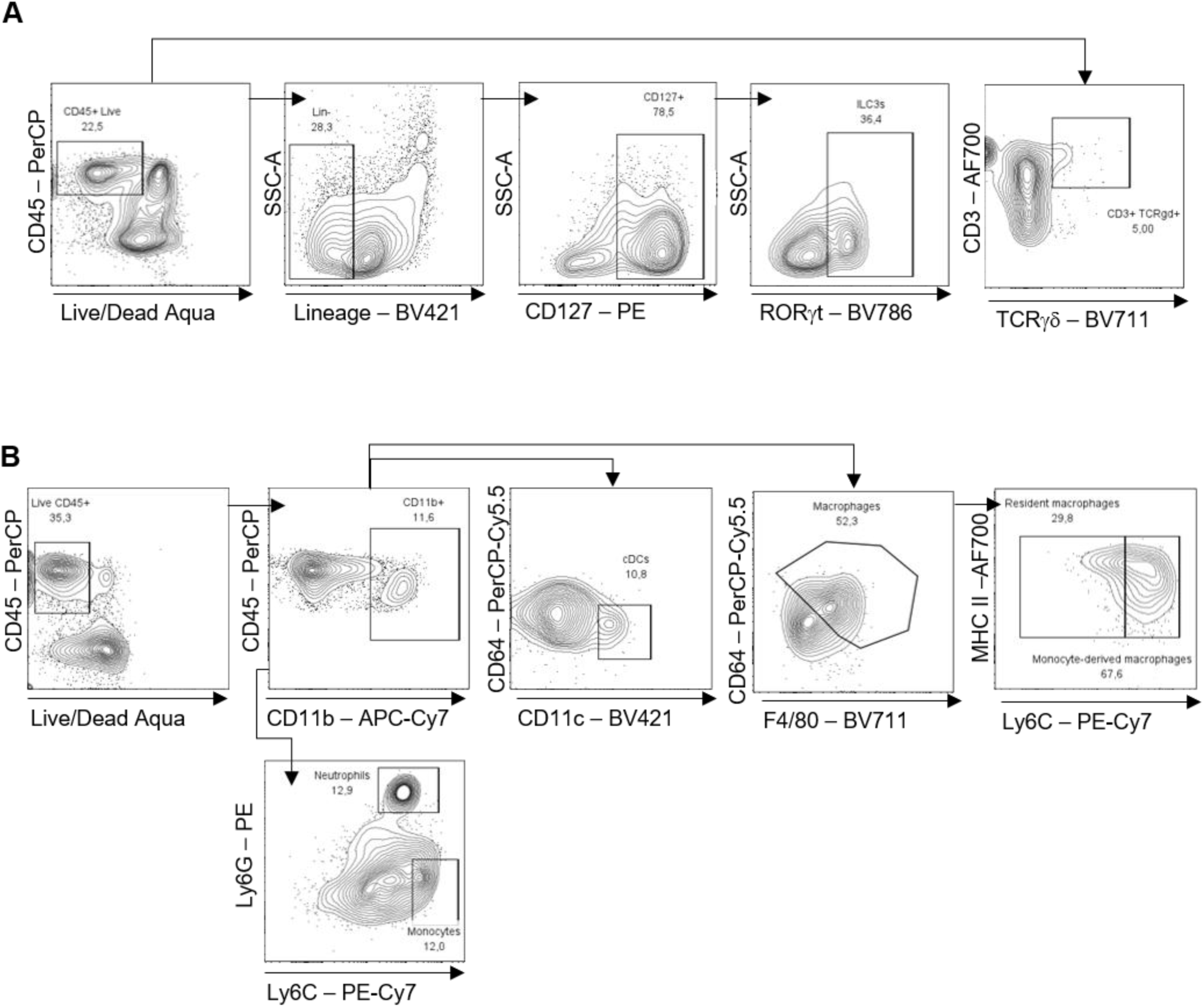
Gating strategy for innate and innate-like immune cells in colonic lamina propria. (A) Sequential identification of live CD45⁺ lineage-negative CD127⁺RORγt⁺ ILC3s and CD3⁺TCRγδ⁺ cells. (B) Sequential myeloid gating using CD11b, CD11c, CD64, F4/80, MHC II, Ly6C, and Ly6G to identify conventional dendritic cells, resident and monocyte-derived macrophages, monocytes, and neutrophils. The gated populations were used to calculate the composition shown in Fig. 2L.

**Figure S4.**
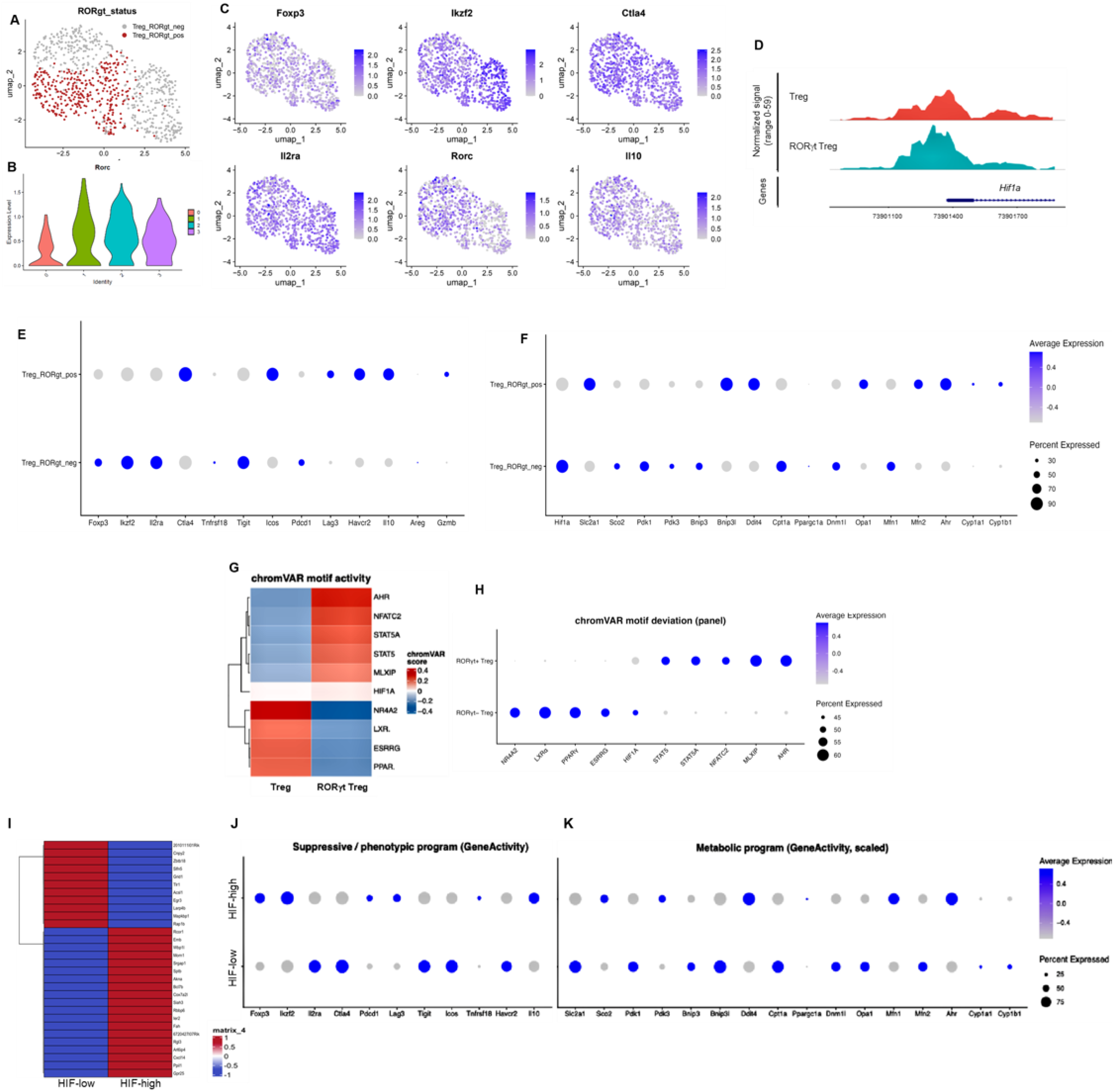
Mouse colonic Treg scATAC-seq nominates regulatory and mitochondrial outputs for functional testing. Reanalysis of mouse colonic Treg scATAC-seq data (GSM6697676). (A) UMAP showing RORγt⁺ and RORγt⁻ Treg. (B) Rorc gene-activity distribution by cluster. (C) Gene-activity feature plots for Foxp3, Ikzf2, Ctla4, Il2ra, Rorc, and Il10. (D) Aggregated accessibility at the Hif1a locus in RORγt⁺ and RORγt⁻ Treg. (E,F) Dot plots of suppressive/phenotypic (E) and metabolic (F) gene activity in RORγt⁺ and RORγt⁻ Treg. (G) Heatmap of chromVAR motif-deviation scores. (H) Dot plot of selected motif-deviation scores. (I) Top differentially accessible peaks between HIF1A motif-low and motif-high RORγt⁺ Treg. (J,K) Suppressive/phenotypic (J) and metabolic (K) gene-activity programs in HIF1A motif-high and motif-low RORγt⁺ Treg. Motif-activity labels denote relative chromVAR deviation and do not represent measured HIF-1α protein abundance or expression.

**Figure S5.**
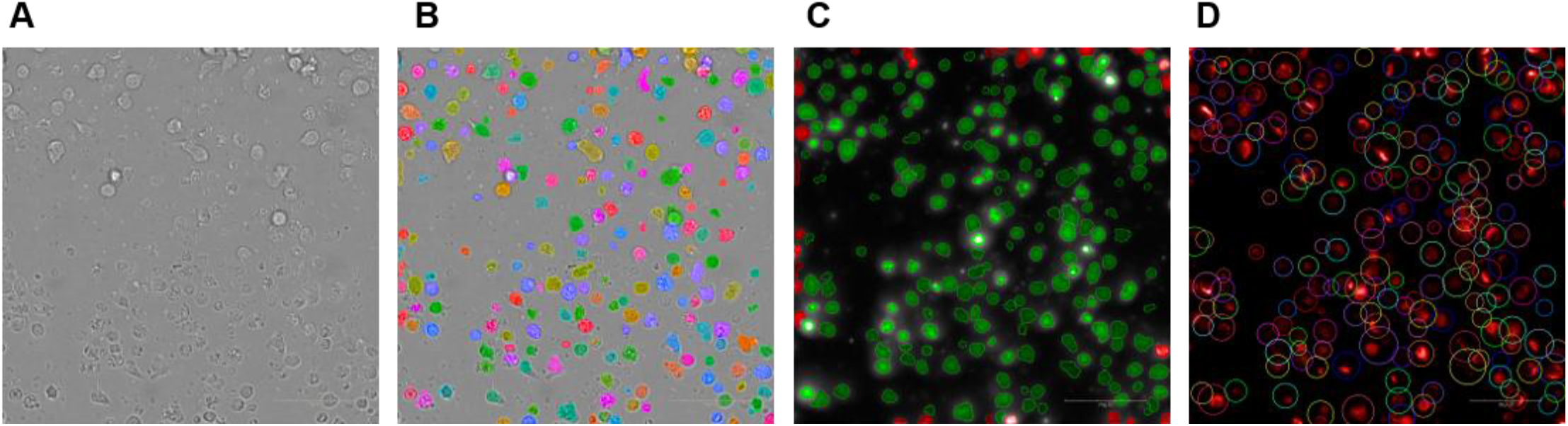
Mask-based workflow for confocal image segmentation and mitochondrial-signal quantification. (A) Representative transmitted-light image of *in vitro*–differentiated RORγt⁺ Treg. (B) Primary segmentation of individual cellular objects, displayed as distinct pseudocolored masks over the transmitted-light image. (C) Detection of SYBR Green–positive objects and definition of the corresponding cell-associated analysis regions, with MitoTracker Red fluorescence retained within the selected objects. (D) Final per-cell regions of interest, displayed as pseudocolored contours over the merged SYBR Green and MitoTracker Red channels. These masks were used to assign mitochondrial fluorescence to individual cells and calculate the mean MitoTracker Red fluorescence intensity per selected cell. Identical acquisition and segmentation settings were applied to WT and ΔHif1a cultures. Scale bars, 50 μm.

**Figure S6.**
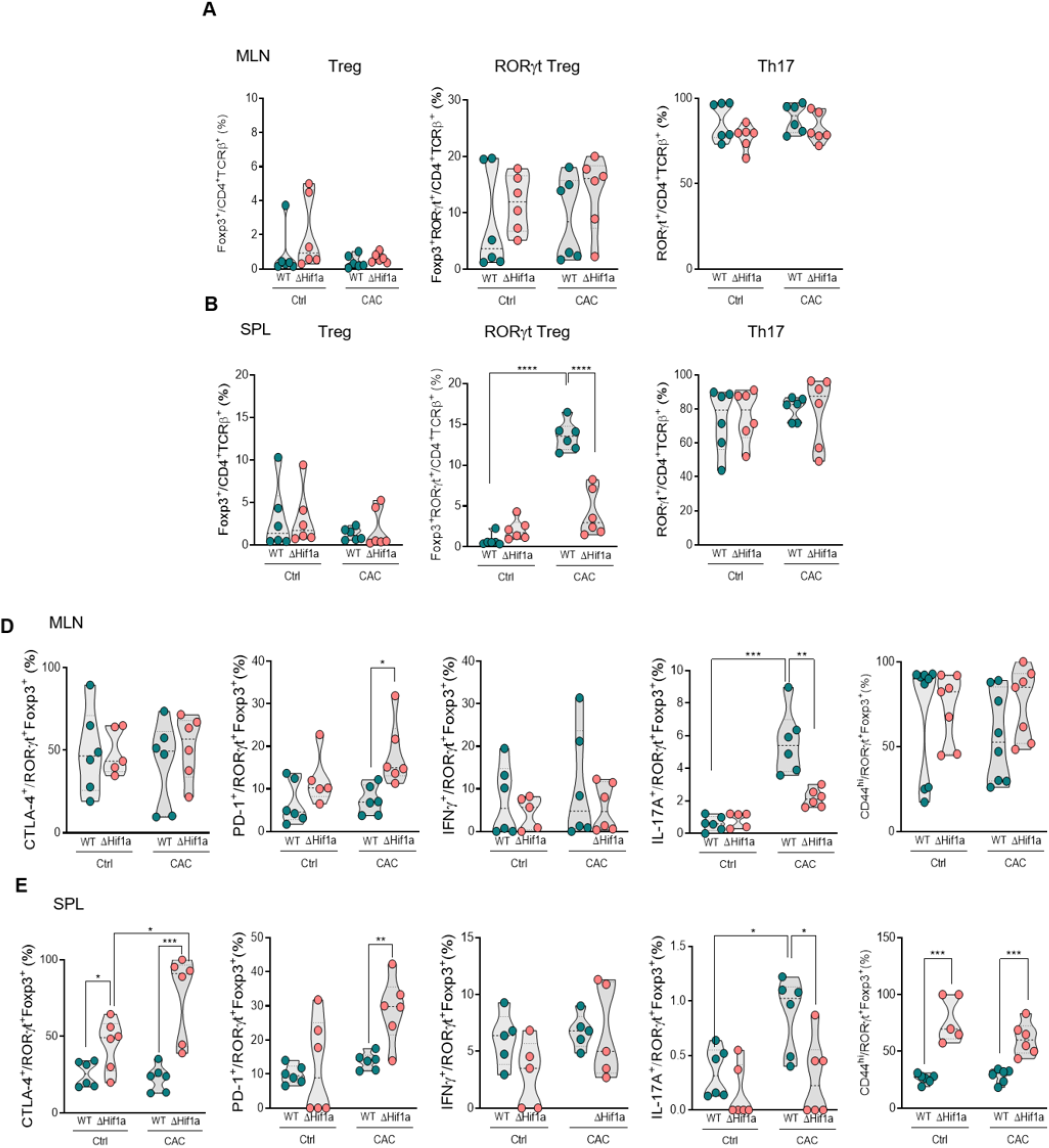
Distribution and functional phenotype of RORγt⁺ Treg in secondary lymphoid organs during CAC. (A,B) Frequencies of Foxp3⁺ Treg, Foxp3⁺RORγt⁺ Treg, and Th17 cells in MLN (A) and spleen (B) from WT and ΔHif1a mice under control or CAC conditions. (D,E) CTLA-4, PD-1, IFN-γ, IL-17A, and CD44 within RORγt⁺ Treg from MLN (D) and spleen (E). Points represent mice; data are mean ± SEM.

**Figure S7.**
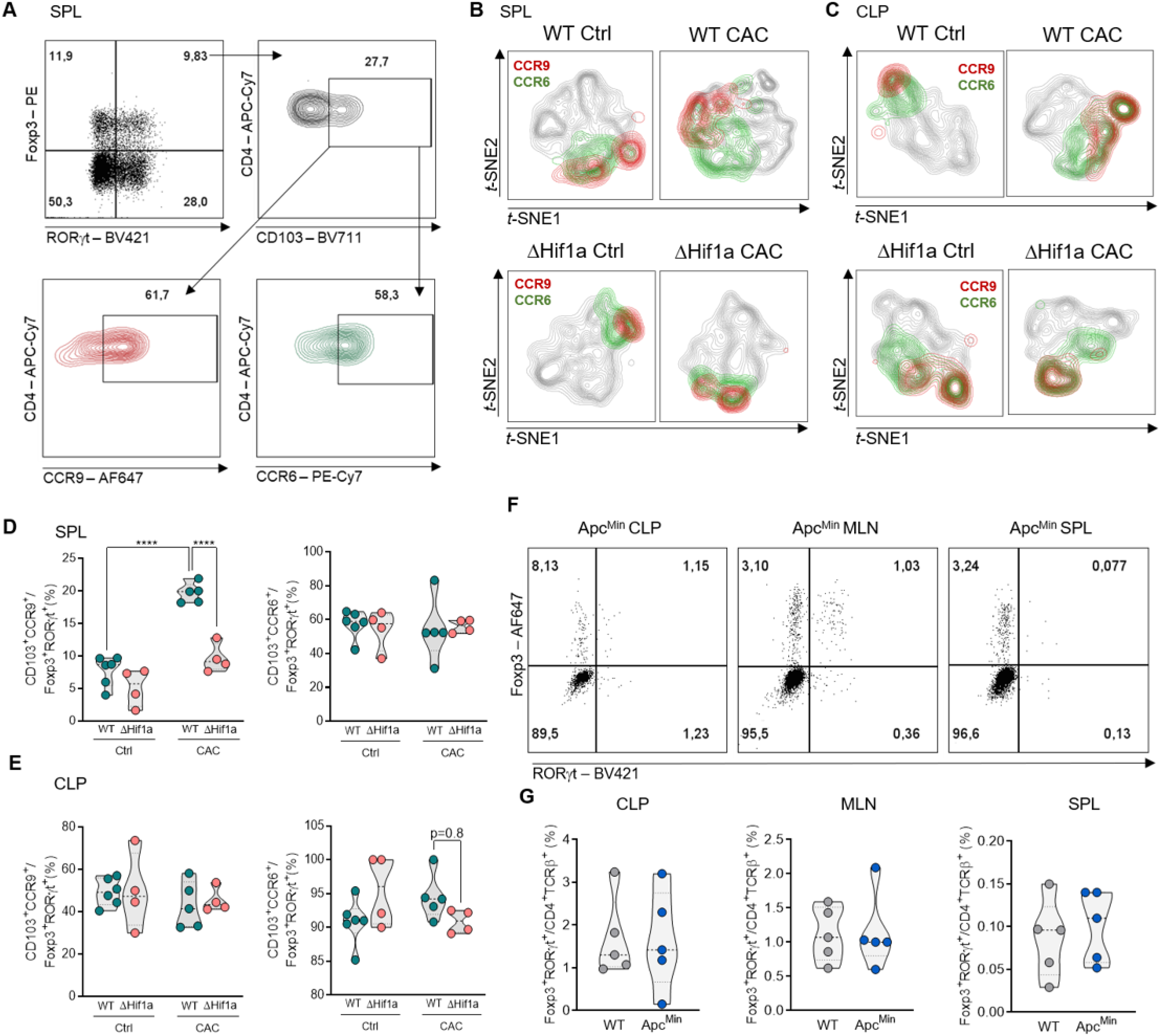
Selective regulation of the CD103⁺CCR9⁺ RORγt⁺ Treg compartment during CAC and analysis of ApcMin mice. (A) Gating of CD103⁺CCR9⁺ and CD103⁺CCR6⁺ cells within splenic Foxp3⁺RORγt⁺ Treg. (B,C) t-SNE maps showing CCR9⁺ and CCR6⁺ cells in spleen (B) and CLP (C) from WT and ΔHif1a mice under control or CAC conditions. (D,E) Frequencies of CD103⁺CCR9⁺ and CD103⁺CCR6⁺ RORγt⁺ Treg in spleen (D) and CLP (E). (F) Representative Foxp3/RORγt plots from CLP, MLN, and spleen of Apc^Min^ mice. (G) Frequencies of Foxp3⁺RORγt⁺ Treg in WT and Apc^Min^ tissues. Points represent mice; data are mean ± SEM.

